# NODAL/TGFβ signalling mediates the self-sustained stemness induced by *PIK3CA*^*H1047R*^ homozygosity in pluripotent stem cells

**DOI:** 10.1101/753830

**Authors:** Ralitsa R. Madsen, James Longden, Rachel G. Knox, Xavier Robin, Franziska Völlmy, Kenneth G. Macleod, Larissa S. Moniz, Neil O. Carragher, Rune Linding, Bart Vanhaesebroeck, Robert K. Semple

## Abstract

Activating *PIK3CA* mutations are known “drivers” of human cancer and developmental overgrowth syndromes. We recently demonstrated that the “hotspot” *PIK3CA*^*H1047R*^ variant exerts unexpected allele dose-dependent effects on stemness in human pluripotent stem cells (hPSCs). In the present study, we combine high-depth transcriptomics, total proteomics and reverse-phase protein arrays to reveal potentially disease-related alterations in heterozygous cells, and to assess the contribution of activated TGFβ signalling to the stemness phenotype of *PIK3CA*^*H1047R*^ homozygous cells. We demonstrate signalling rewiring as a function of oncogenic PI3K signalling dose, and provide experimental evidence that self-sustained stemness is causally related to enhanced autocrine NODAL/TGFβ signalling. A significant transcriptomic signature of TGFβ pathway activation in *PIK3CA*^*H1047R*^ heterozygous was observed but was modest and was not associated with the stemness phenotype seen in homozygous mutants. Notably, the stemness gene expression in *PIK3CA*^*H1047R*^ homozygous iPSCs was reversed by pharmacological inhibition of TGFβ signalling, but not by pharmacological PI3Kα pathway inhibition. Altogether, this provides the first in-depth analysis of PI3K signalling in human pluripotent stem cells and directly links dose-dependent PI3K activation to developmental NODAL/TGFβ signalling.

## Introduction

Class IA phosphoinositide 3-kinases (PI3Ks) are evolutionarily conserved enzymes that catalyse formation of the membrane-bound second messenger phosphatidylinositol-3,4,5-trisphosphate (PIP_3_). PI3Ks are activated downstream of receptor tyrosine kinases, with the ensuing increase in PIP_3_ and its derivative PI(3,4)P_2_ triggering a widespread signalling network, best known for the activation of the serine/threonine kinases AKT and mTORC1. PI3K activation promotes cell survival, glucose uptake, anabolic metabolism, cell proliferation and cell migration (1). Among the class IA PI3K isoforms (PI3Kα, PI3Kβ, PI3Kδ), the ubiquitously-expressed PI3Kα (encoded by the *PIK3CA* gene in humans), is the main regulator of organismal growth, development and survival (2).

Activating mutations in *PIK3CA* are among the most common somatic point mutations in cancer, together with inactivation or loss of the tumour suppressor *PTEN* (a negative regulator of PI3K) (3–5). The same mutations in *PIK3CA*, when acquired postzygotically during development, also cause a range of largely benign overgrowth disorders, for which the term *PIK3CA*-related overgrowth spectrum (PROS) has been coined (6). Motivated by the need to understand the role of PI3K signalling in a human developmental context, we previously generated an allelic series of human induced pluripotent stem cells (iPSCs) with heterozygous or homozygous expression of the activating mutation *PIK3CA*^*H1047R*^, the most commonly observed *PIK3CA* mutation in both cancer and PROS (7). Despite the severe developmental disorders caused by heterozygosity for *PIK3CA*^*H1047R*^ in humans *in vivo*, we found little discernible effect on germ layer specification from heterozygous iPSCs. In sharp contrast, homozygosity for *PIK3CA*^*H1047R*^ led to self-sustained stemness and resistance to spontaneous differentiation *in vitro* and *in vivo* (7). This suggested a previously unappreciated quantitative relationship between the strength of PI3K signalling and the gene regulatory network (GRN) in pluripotent stem cells.

The core pluripotency GRN features a feedforward, autoregulatory circuit comprising three transcription factors, namely SRY box 2 (SOX2), Octamer-binding transcription factor 3/4 (OCT3/4; encoded by POU5F1), and the homeobox transcription factor NANOG (8–10). SOX2 helps sustain OCT3/4 expression, which is required for establishment and maintenance of the pluripotent state (11). However, even modest overexpression of OCT3/4 destabilises the pluripotency network and triggers differentiation (12, 13). In contrast, NANOG, while dispensable for maintenance of pluripotency (14), stabilises the pluripotency gene regulatory network. Overexpression of NANOG by as little as 1.5-fold leads to sustained self-renewal (or “stemness”) of murine and human PSCs (15–18). In hPSCs, NANOG expression is activated by the transcription factors SMAD2/3 (19), which in turn are activated by receptors binding TGFβ, Activin or NODAL (20). Overexpression of NODAL thus results in self-sustained stemness of hPSCs even in differentiation-promoting conditions (21, 22).

Given the unexpected and surprisingly mild phenotype caused by heterozygous *PIK3CA*^*H1047R*^ expression in iPSCs, we reasoned that more sensitive assays would allow us to discern small but disease-relevant alterations in these cells. Thus, in this study, we first applied high depth transcriptomics, and proteomics to seek evidence of disease-related phenotypes in heterozygous cells, and to investigate how high-dose PI3K signalling leads to self-sustained stemness in homozygous *PIK3CA*^*H1047R*^ iPSCs. We demonstrate that heterozygous cells do exhibit significant transcriptomic changes, although these are a weak echo of the widespread changes seen in homozygous cells. The mild transcriptional consequences of heterozygous expression of disease-relevant *PIK3CA* mutations were also validated in additional model systems and contrast with previous findings of major transcriptional rewiring in immortalised, non-transformed breast epithelial cells (23, 24). We demonstrate that the stemness phenotype of *PIK3CA*^*H1047R/H1047R*^ iPSCs is maintained by self-sustained NODAL/TGFβ signalling, in line with increased *PIK3CA*-mediated *NODAL* expression, and that it is not reversible by PI3Kα-specific inhibition. This work provides the first in-depth characterisation of dose-dependent PI3K signalling effects in hPSCs and establishes dose-dependent PI3Kα-induced NODAL/TGFβ signalling as the main mechanism for self-sustained stemness in homozygous *PIK3CA*^*H1047R*^ iPSCs. We discuss the implications of our findings for understanding developmental disorders and cancers driven by genetic PI3K activation.

## Results

### A sharp PI3K activity threshold determines gene expression changes in *PIK3CA*^*H1047R*^ iPSCs

We previously generated isogenic human iPSCs with heterozygous or homozygous knock-in of the “hotspot” *PIK3CA*^*H1047R*^ mutation. Surprisingly, heterozygous cells showed few phenotypic changes and differentially expressed protein-coding transcripts. In contrast, homozygous *PIK3CA*^*H1047R/H1047R*^ cells exhibited marked morphological changes and altered gene expression, with strong enrichment for cancer-associated pathways (7).

To substantiate the apparent PI3K activity threshold manifest in *PIK3CA*^*H1047R*^-driven gene expression changes, and to look for further disease-related changes in heterozygous cells, we undertook RNA sequencing at substantially greater depth, also increasing the sample size to four independently-derived, previously unstudied iPSC cultures for each *PIK3CA* genotype. As before, homozygous mutant cells clearly separated from heterozygous and wild-type cells, which overlapped on multidimensional scaling (**Fig. 1A**), but we now detected a reduction in the levels of 451 transcripts and an increase in the levels of 710 transcripts in *PIK3CA*^*WT/H1047R*^ iPSCs (**Fig. 1B**). This dropped to 149 and 343 transcripts, respectively, after applying a fold-change cut-off of 1.3 (**Fig. 1B and Dataset S1**), indicative of the small magnitude of many expression changes in heterozygous mutants (**Fig. S1A**). Use of the same cut-off of 1.3, in sharp distinction, yielded 2873 and 2771 transcripts of decreased or increased abundance, respectively, in homozygous iPSC mutants (**Fig. 1B and Dataset S2**). Not only was the number of gene expression changes higher by an order of magnitude in homozygous cells, but many expression changes were large compared to wild-type controls (**Fig. S1A**). The magnitudes of gene expression changes in *PIK3CA*^*H1047R/H1047R*^ cells correlated strongly with our previous findings (Spearman’s rho = 0.74, p < 2e-16) (**Fig. S1B**), whereas correlation was low (Spearman’s rho = 0.1, p < 2e-16) for *PIK3CA*^*WT/H1047R*^ iPSCs (**Fig. S1C**).

**Figure 1.**
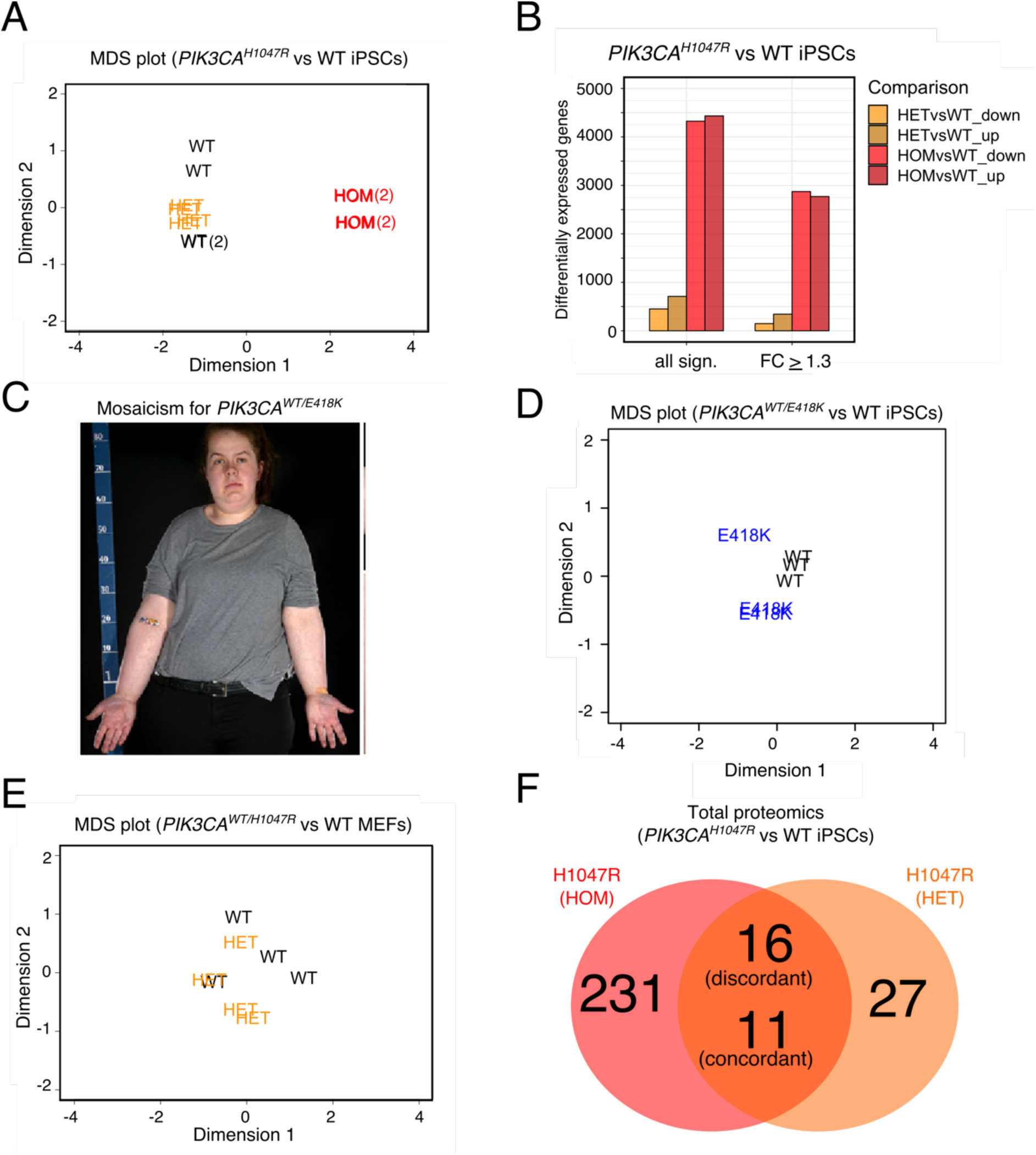
Transcriptomic and proteomic analyses of human and mouse cell lines with endogenous expression of oncogenic *PIK3CA*. **(A)** Multidimensional scaling (MDS) plot of the transcriptomes of wild-type (WT), *PIK3CA*^*WT/H1047R*^ (HET) and *PIK3CA*^*H1047R/H1047R*^ (HOM) human iPSCs. The numbers in brackets indicate the presence of two closely overlapping samples. **(B)** The number of differentially expressed genes in iPSCs heterozygous or homozygous for *PIK3CA*^*H1047R*^ before and after application of an absolute fold-change cut-off ≥ 1.3 (FDR ≤ 0.05, Benjamini-Hochberg). The data are based on four iPSCs cultures from minimum two clones per genotype. See also Fig. S1. **(C)** Woman with asymmetric overgrowth caused by mosaicism for cells with heterozygous expression of *PIK3CA*^*E418K*^. Skin biopsies obtained from unaffected and affected tissues were used to obtain otherwise isogenic dermal fibroblasts for subsequent reprogramming into iPSCs. This image was reproduced from Ref. (25) **(D)** MDS plot of the transcriptomes of wild-type (WT) and *PIK3CA*^*WT/E418K*^ iPSCs (based on 3 independent mutant clones and 3 wild-type cultures from 2 independent clones). **(E)** MDS plot of the transcriptomes of wild-type (WT) and *PIK3CA*^*WT/H1047R*^ (HET) mouse embryonic fibroblasts (MEFs) following 48 h of mutant induction (N = 4 independent clones per genotype). **(F)** Venn diagram showing the number of differentially expressed proteins in *PIK3CA*^*H1047R/H1047R*^ (HOM) and *PIK3CA*^*WT/H1047R*^ (HET) iPSCs relative to wild-type controls, profiled by label-free total proteomics on three clones per genotype. An absolute fold-change and z-score ≥ 1.2 were used to classify proteins as differentially expressed. The number of discordant and concordant changes in the expression of total proteins detected in both comparisons are indicated. See also Fig. S2.

Given prior reports that *PIK3CA*^*H1047R*^ heterozygosity in breast epithelial cells extensively remodels gene expression (23, 24), we undertook further transcriptional profiling in two unrelated cellular models of genetic *PIK3CA* activation. First, we examined iPSCs derived from a woman with clinically obvious but mild PROS due to mosaicism for *PIK3CA*^*E418K*^ (**Fig. 1C**) (25). Heterozygous iPSCs were compared to wild-type lines established simultaneously from dermal fibroblasts from the same skin biopsy, which is possible due to genetic mosaicism of the sampled skin. Like *PIK3CA*^*WT/H1047R*^ iPSCs, *PIK3CA*^*WT/E418K*^ iPSCs closely clustered with isogenic wild-type controls on multidimensional scaling (MDS) plotting (**Fig. 1D**), with only 30 differentially expressed genes (**Dataset S3**). We also studied previously reported *Pik3ca*^*WT/H1047R*^ mouse embryonic fibroblasts (MEFs) 48 h after *Cre*-mediated *Pik3ca*^*H1047R*^ induction (26). Wild-type and *Pik3ca*^*WT/H1047R*^ MEFs were superimposable on an MDS plot (**Fig. 1E**), with only 192 downregulated and 77 upregulated genes (**Dataset S4**). Our findings suggest that there are *bona fide* transcriptional changes induced by heterozygosity for *PIK3CA*^*H1047R*^, but these are dramatically smaller in number and magnitude than changes induced by homozygosity for *PIK3CA*^*H1047R*^.

To assess whether transcriptional changes observed in iPSCs were mirrored in the proteome, we applied label-free proteomics to the iPSC lines used in our previous study (7). Around 4,600 protein ratios were obtained for both heterozygous *versus* wild-type and homozygous *versus* wild-type iPSC comparisons, as estimated using a novel Bayesian approach based on the Markov Chain Monte Carlo (MCMC) method (27). In contrast to other algorithms, the MCMC method generates an error estimate alongside each protein concentration which permits more confident determination of proteins with the most robust differential expression. The number of differentially-expressed proteins correlated with *PIK3CA*^*H1047R*^ allele dosage, with 54 and 258 differentially expressed proteins in *PIK3CA*^*WT/H1047R*^ and *PIK3CA*^*H1047R/H1047R*^ cells, respectively (**Fig. 1F, Datasets S5 and S6)**. Of these, 27 proteins were differentially expressed in both heterozygous and homozygous *PIK3CA*^*H1047R*^ iPSCs (**Dataset S7**), with 16 changing in opposite directions (**Fig. 1F**). There was a strong correlation between differentially-expressed proteins and corresponding transcripts in *PIK3CA*^*H1047R/H1047R*^ iPSCs (**Fig. S2A, S2B**), but not in heterozygous mutants (**Fig. S2C, S2D**). As for the relatively weak correlation seen between transcriptomic experiments for heterozygous cells, this likely reflects the small magnitude of gene expression changes induced by heterozygous *PIK3CA*^*H1047R*^ (**Fig. 1B, S1C**).

Collectively, these findings corroborate the existence of a threshold of PI3K pathway activity which determines the large majority of gene expression changes in *PIK3CA*^*H1047R/H1047R*^ iPSCs in a near-binary manner. While deeper sequencing did reveal statistically significant gene expression changes in heterozygous iPSCs, and while these changes may contribute to growth-related phenotypes in PROS when sustained across development, effect sizes were modest and more variable. Similar findings in heterozygous MEFs suggest that this may be generalisable to differentiated cell types, irrespective of species. This consolidates the view that only homozygosity for *PIK3CA*^*H1047R*^ results in robust and widespread transcriptional changes in otherwise normal, diploid cells, arguing against a universal “butterfly” effect of heterozygosity suggested based on studies of a genetically abnormal breast epithelial cell line (23, 24).

### *PIK3CA*^*H1047R/H1047R*^ iPSCs show evidence of signalling “rewiring”

We previously demonstrated a graded increase in AKT (S473) phosphorylation across heterozygous and homozygous *PIK3CA*^*H1047R*^ iPSCs (7). To assess in more detail whether the near-binary gene expression difference between heterozygous and homozygous *PIK3CA*^*H1047R*^ cells is underpinned by corresponding differences in indices of PI3K pathway activation, we profiled phosphorylation of a wider repertoire of pathway components using reverse phase phosphoprotein array (RPPA) technology.

Changes in protein phosphorylation were surprisingly modest, with the largest change a two-fold increase in AKT phosphorylation (on S473 and T308) in *PIK3CA*^*H1047R/H1047R*^ cells. Contrasting with the near-binary response seen at the transcriptional level, heterozygous and homozygous *PIK3CA*^*H1047R*^ expression generally produced graded phosphorylation of PI3K pathway components, with slightly higher levels in homozygous iPSCs (**Fig. 2A**). None of the mutant genotypes showed consistently increased phosphorylation of the mTORC1 target P70S6K or its downstream substrate S6 (**Fig. S3A**), perhaps reflecting saturation at this level of the pathway due to other stimuli for mTORC1 in the complete culture medium (e.g. amino acids) (28). When deprived of growth factors for 1 h prior to RPPA profiling, both heterozygous and homozygous mutant did exhibit increased P70S6K phosphorylation, whereas S6 phosphorylation remained similar to wild-type cells (**Fig. 2B**).

**Figure 2.**
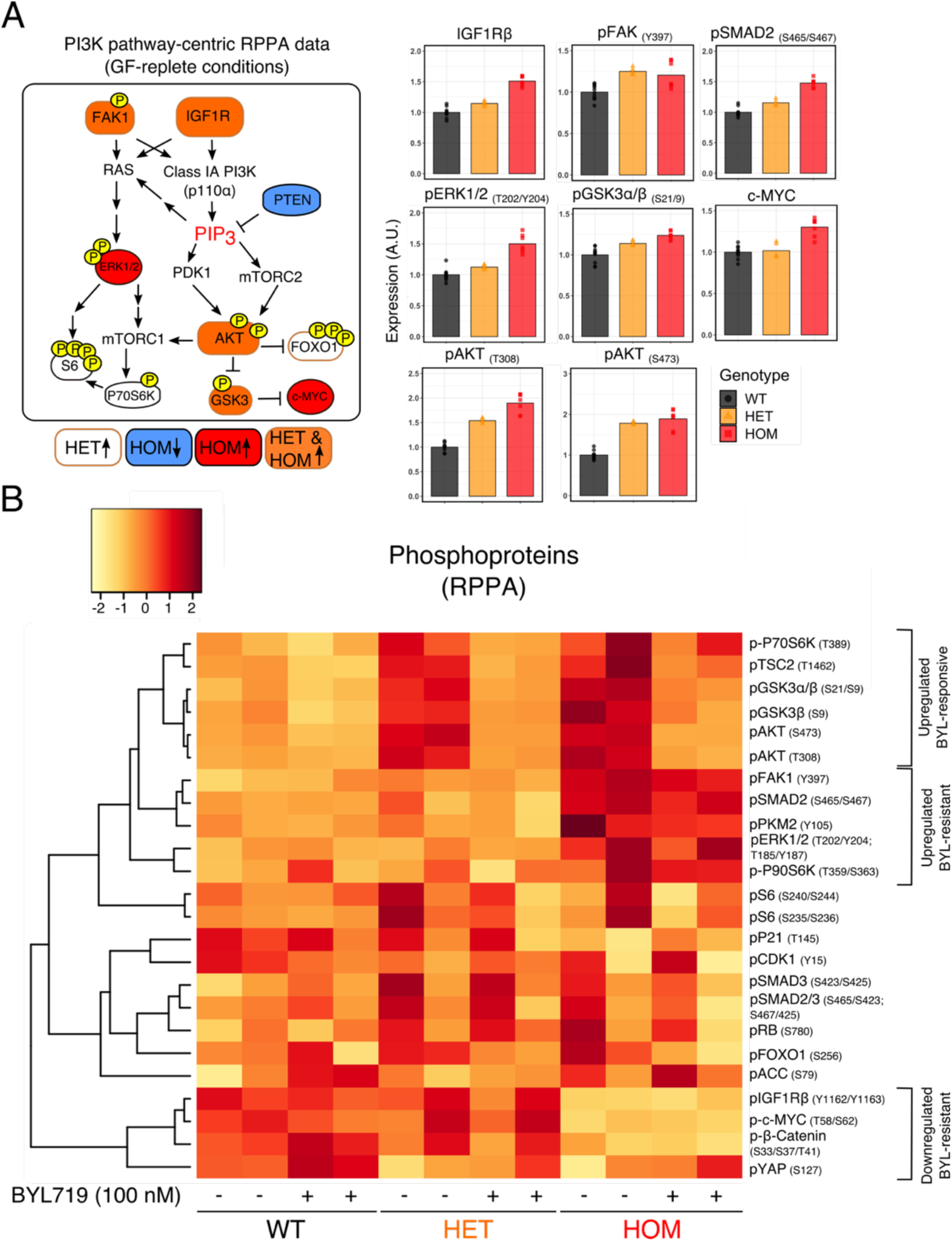
Reverse Phase Protein Array (RPPA) of *PIK3CA*^*WT/H1047R*^ (HET) and *PIK3CA*^*H1047R/H1047R*^ (HOM) human iPSCs. **(A)** Left: Diagram of PI3K pathway-related phosphorylated proteins, with colour code used to signify differentially expressed targets in *PIK3CA*^*H1047R*^ mutant iPSCs *versus* isogenic wild-type controls. Color-coded targets were significant at FDR ≤ 0.05 (Benjamini-Hochberg). Right: Barplots show representative examples of differentially expressed protein targets, revealing relatively modest quantitative changes. Phosphorylated proteins were normalised to the corresponding total protein when available. The data are based on 10 wild-type cultures (3 clones), 5 *PIK3CA*^*WT/H1047R*^ cultures (3 clones) and 7 *PIK3CA*^*H1047R/H1047R*^ cultures (2 clones) as indicated. See also Fig. S3A. **(B)** Unsupervised hierarchical clustering based on target-wise correlations of RPPA data from wild-type (WT), *PIK3CA*^*WT/H1047R*^ (HET) and *PIK3CA*^*H1047R/H1047R*^ (HOM) iPSCs following short-term growth factor removal (1 h), +/- 100 nM BYL719 (PI3Kα inhibitor) for 24 h. The data are from two independent experiments, each performed using independent clones. For each row, the colours correspond to Fast Green-normalised expression values in units of standard deviation (z-score) from the mean (centred at 0) across all samples (columns). Groups of phosphorylated proteins exhibiting a consistent expression pattern in BYL719-treated *PIK3CA*^*H1047R/H1047R*^ iPSCs are specified. See also Fig. S3B.

Inhibition of PI3Kα activity with the PI3Kα-selective inhibitor BYL719 for 24 h fully reversed canonical PI3K signalling-related changes in phosphorylation of downstream proteins including AKT, GSK3, FOXO1, TSC2 and P70S6K (**Fig. 2B**). Consistent with these signalling changes, we previously showed that the same dose of BYL719 (100 nM) abolishes the increased tolerance to growth factor deprivation-induced death conferred by heterozygous or homozygous *PIK3CA*^*H1047R*^ in iPSCs (7). Despite its effects on the primary PI3K signalling cascade, PI3Kα inhibition failed to reverse other changes observed in *PIK3CA*^*H1047R/H1047R*^ iPSCs, including increased phosphorylation of SMAD2 and ERK1/2 and increased expression of c-MYC and IGF1R (**Fig. 2B, Fig. S3B**). This suggests signalling rewiring in *PIK3CA*^*H1047R/H1047R*^ iPSCs that is partially resistant to relatively short-term inhibition of the inducing stimulus.

### Pathway and network analyses implicate TGFβ signalling in *PIK3CA*^*H1047R*^ dose-dependent stemness

Pathway and network analyses were next applied to proteomic and transcriptomic data to identify candidate mechanism(s) mediating *PIK3CA*^*H1047R*^ dose-dependent stemness. Consistent with our previous study (7), TGFβ1 was again the most significant predicted upstream activator according to Ingenuity^®^ Pathway Analysis (IPA) of the top 2000 upregulated and top 2000 downregulated transcripts in *PIK3CA*^*H1047R/H1047R*^ iPSCs (**Fig. 3A**). TGFβ1 was also the most significant upstream activator predicted by analysis of *PIK3CA*^*H1047R/H1047R*^ proteomic data (**Fig. 3B**). This strongly suggests activation of the TGFβ pathway in homozygous *PIK3CA*^*H1047R*^ iPSCs.

**Figure 3.**
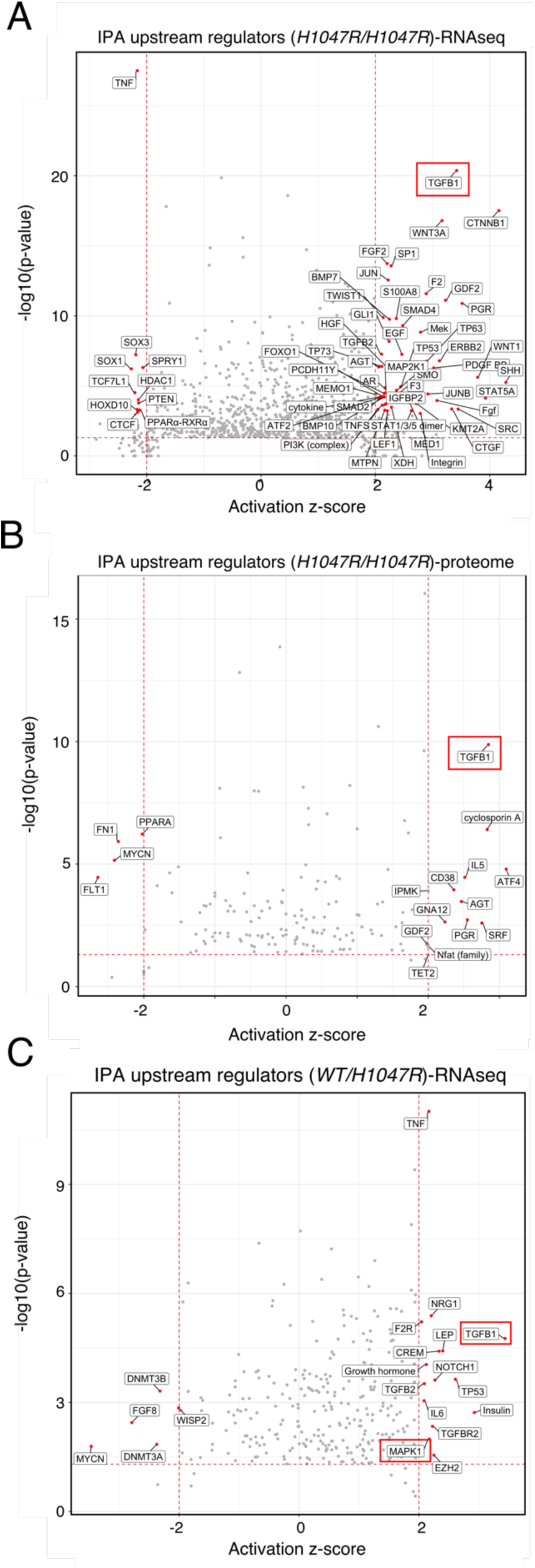
Ingenuity® pathway analyses (IPA) predict activation of TGFβ signalling in heterozygous and homozygous *PIK3CA*^*H1047R*^ iPSCs. **(A)** IPA of upstream regulators using the list of top 2000 upregulated and top 2000 downregulated mRNA transcripts in *PIK3CA*^*H1047R/H1047R*^ iPSCs (for RNAseq details, see Fig. 1B). Red points signify transcripts with absolute predicted activation z-score > 2 and overlap P-value < 0.001 (Fisher’s Exact Test). The red rectangle highlights the most significant upstream regulator, TGFβ1. **(B)** As in (A) but using the list of differentially-expressed proteins identified by total proteomics and red-colouring targets with predicted activation z-score > 2 and overlap P-value < 0.05 (Fisher’s Exact Test). **(C)** As in (A) but using the list of differentially expressed total proteins in *PIK3CA*^*WT/H1047R*^ iPSCs and red-colouring upstream regulators with absolute predicted bias-corrected z score > 2 and overlap P-value < 0.05 (Fisher’s Exact Test). Red rectangles highlight the two upstream regulators (TGFβ1 and MAPK1) with absolute predicted bias-corrected z score > 2 that remained significant (overlap P-value < 0.05) when the analysis was repeated using the list of shared and concordant differentially expressed genes (N = 180) in heterozygous and homozygous *PIK3CA*^*H1047R*^ iPSCs *vs* wild-type controls.

Although *PIK3CA*^*WT/H1047R*^ iPSCs showed around 10-fold fewer differentially expressed genes than homozygous iPSC cells, IPA in heterozygous iPSCs also revealed multiple TGFβ pathway-related stimuli among predicted upstream activators (**Fig. 3C**). Moreover, TGFβ1 was predicted as one of only two significant upstream activators when analysis was performed on genes concordantly differentially expressed (N = 180) in *PIK3CA*^*H1047R*^ mutant iPSCs *versus* wild-type controls (**Fig. 3C and Dataset S8**).

The other significant upstream regulator common to heterozygous and homozygous *PIK3CA*^*H1047R*^ was MAPK1 (also known as ERK2), consistent with RPPA findings and immunoblot evidence of increased ERK kinase phosphorylation in *PIK3CA*^*H1047R*^ mutant iPSCs (Ref. (7), **Fig. 2A and Fig. S3A**). The significance of predicted TGFβ activation in heterozygous *PIK3CA*^*H1047R*^ iPSCs (overlap p-value = 1.7e-05) was much lower than in homozygous (overlap p-value = 4.3e-21) mutants. This points towards a critical role for the TGFβ pathway in mediating the allele dose-dependent effect of *PIK3CA*^*H1047R*^ in human iPSCs.

To complement IPA analysis, which is based on highly curated, proprietary datasets, we undertook non-hypothesis-based Weighted Gene Correlation Network Analysis (WGCNA) – a network-based data reduction method that seeks to determine gene correlation patterns across multiple samples, irrespective of the function of individual genes (29). Using all transcripts expressed in wild-type, heterozygous and homozygous *PIK3CA*^*H1047R*^ iPSCs (**Fig. 4A**), this analysis returned 43 modules (or clusters) of highly interconnected genes (**Fig. 4B**). Of the two modules with the highest correlation with the homozygous trait, one showed enrichment for several KEGG pathway terms relevant to stemness of *PIK3CA*^*H1047R/H1047R*^ iPSCs, notably including “Signalling pathways regulating pluripotency in stem cells” (**Fig. 4C**).

**Figure 4.**
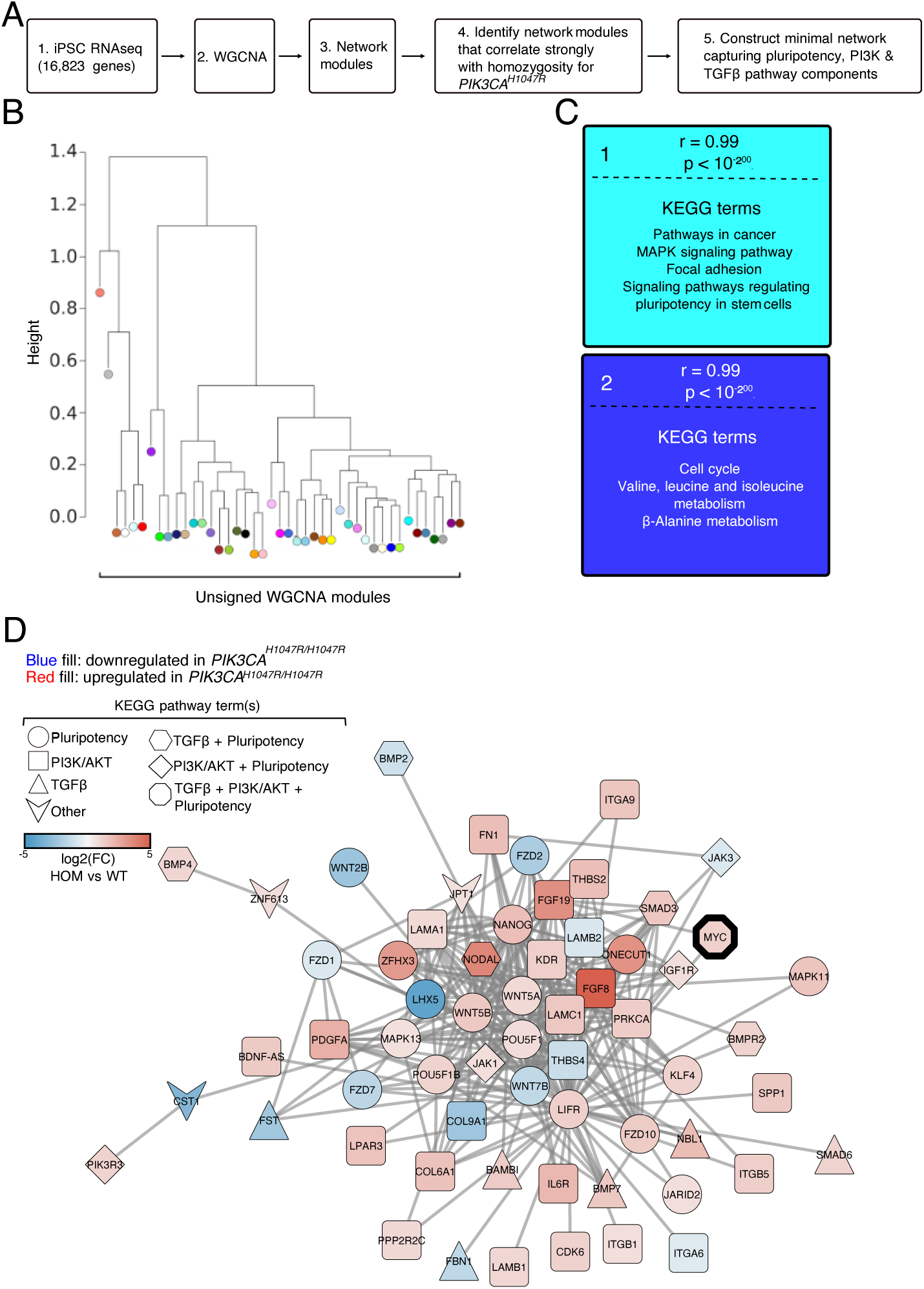
Weighted gene correlation network analysis (WGCNA) identifies links among pluripotency components, TGFβ and PI3K signalling. **(A)** Schematic of the WGCNA workflow and subsequent data selection for visualisation. (**B)** Unsigned WGCNA modules identified using the list of transcripts expressed in wild-type, *PIK3CA*^*WT/H1047R*^ and *PIK3CA*^*H1047R/H1047R*^ iPSCs (for RNAseq details, see Fig. 1B). **(C)** The two network modules with genes whose module membership correlated strongest with differential expression in homozygous *PIK3CA*^*H1047R*^ iPSCs. The colour of each module corresponds to its colour in the module dendrogram in (B). Representative KEGG pathways with significant enrichment in each gene network module are listed (hypergeometric test with two-sided uncorrected P < 0.05). **(D)** The minimal network connecting KEGG pluripotency, PI3K/AKT and TGFβ pathway components within the turquoise gene network module. Fill colour and shape are used to specify direction of differential mRNA expression in *PIK3CA*^*H1047R/H1047R*^ iPSCs and pathway membership, respectively. Fill colour saturation represents gene expression fold-change (FC; log2) in *PIK3CA*^*H1047R/H1047R*^ (HOM) *vs* wild-type (WT) iPSCs. MYC is highlighted as the only network component intersecting all three KEGG pathways, suggesting it may comprise a key mechanistic link in the observed phenotype.

Given prior evidence of strong activation of TGFβ signalling in homozygous mutant cells, we next constructed the minimal network of differentially expressed genes in *PIK3CA*^*H1047R/H1047R*^ iPSCs that linked pluripotency, PI3K and TGFβ signalling pathways (**Fig. 4D**). This approach allowed us to navigate the signalling rewiring and to link strong PI3K pathway activation, stemness and TGFβ signalling in an unbiased manner. Indeed, the resulting network exhibited high interconnectivity, with multiple shared nodes across all three pathways, suggesting close crosstalk between PI3K and TGFβ signalling in stemness regulation. That most nodes represented genes with increased expression in homozygous mutants strengthens the notion that strong oncogenic PI3Kα activation stabilises the pluripotency network in human iPSCs. The MYC oncogene stood out as the only network node intersecting with all three signalling pathways, suggesting it may comprise a key mechanistic link in the observed phenotype.

### Inhibition of TGFβ signalling destabilises the pluripotency gene network in *PIK3CA*^*H1047R/H1047R*^ iPSCs

TGFβ signalling plays a critical role in pluripotency regulation (19, 22, 30), and a differentiation-resistant phenotype has been reported in *NODAL*-overexpressing iPSCs (21). Together with increased *NODAL* expression in homozygous *PIK3CA*^*H1047R*^ iPSCs and computational identification of enhanced TGFβ pathway activity in PI3K-driven “constitutive” stemness (Ref. (7) and current study), this led us to hypothesise that strong PI3Kα-dependent induction of *NODAL* underlies establishment of the differentiation-resistant phenotype of homozygous *PIK3CA*^*H1047R*^ iPSCs. Specifically, we hypothesised that autocrine NODAL enhances TGFβ signalling in *PIK3CA*^*H1047R/H1047R*^ iPSCs, with resulting increased *NANOG* expression “locking” the cells in perpetual stemness (19).

Testing this hypothesis in iPSCs is challenging for biological and technical reasons, including lack of specific pharmacological inhibitors of NODAL, and difficulty in detecting subtle early phenotypic consequences of partial destabilisation of the iPSC pluripotency gene regulatory network. Moreover, the widely adopted maintenance medium and coating substrate we used for cell culture both contain TGFβ ligands (31, 32), which may mask effects of *NODAL* repression by PI3Kα-specific inhibition. We previously found that treatment of *PIK3CA*^*H1047R/H1047R*^ iPSCs in this ‘complete’ maintenance medium with 500 nM BYL719 reduces *NODAL* mRNA expression within 24 h, but has no discernible effect on increased *NANOG* mRNA levels (7).

To minimise confounding effects of exogenous TGFβ ligands, we prepared medium with and without recombinant NODAL supplementation, and assessed expression of *NODAL* and *NANOG* as surrogate markers of stemness over 72 h of culture. We also reduced the BYL719 concentration to 250 nM given increased iPSC toxicity observed with 500 nM BYL719 (7); and pilot experiments (not shown) in which 24 h treatment with 250 nM but not 100 nM BYL719 in complete medium reduced *NODAL* mRNA expression in *PIK3CA*^*H1047R/H1047R*^ iPSC clones. Within 48 h, exclusion of NODAL from the medium resulted in the expected downregulation of *NODAL* and *NANOG* expression in wild-type iPSCs, and this was greater still at 72 h (**Fig. 5 and Fig. S4A**). In *PIK3CA*^*H1047R/H1047R*^ iPSCs, however, *NODAL* removal had no effect on the increased *NODAL* and *NANOG* expression (**Fig. 5 and Fig. S4A**), in line with a self-sustained stemness phenotype. Exposure of NODAL-free *PIK3CA*^*H1047R/H1047R*^ cultures to 250 nM BYL719 had a visible colony growth-inhibitory effect **(Fig. S5**) and decreased *NODAL* expression within 24 h, and this continued to decrease subsequently (**Fig. 5**). This is consistent with NODAL’s known ability to control its own expression through a feed-forward loop (33). Despite a 55% reduction in *NODAL* mRNA after 72 h, however, little effect on *NANOG* expression was seen (**Fig. 5**). This may reflect the short time course studied (to avoid confounding effect of passaging), or the exquisite sensitivity of iPSCs to residual upregulation of *NODAL* in homozygous *PIK3CA*^*H1047R*^ iPSCs. This may be compounded by residual low levels of TGFβ-like ligands in the coating substrate, or possibly by increased expression of two other TGFβ superfamily ligands, *GDF3* and *TGFB2*, observed in homozygous mutant cells (**Dataset S2**).

**Figure 5.**
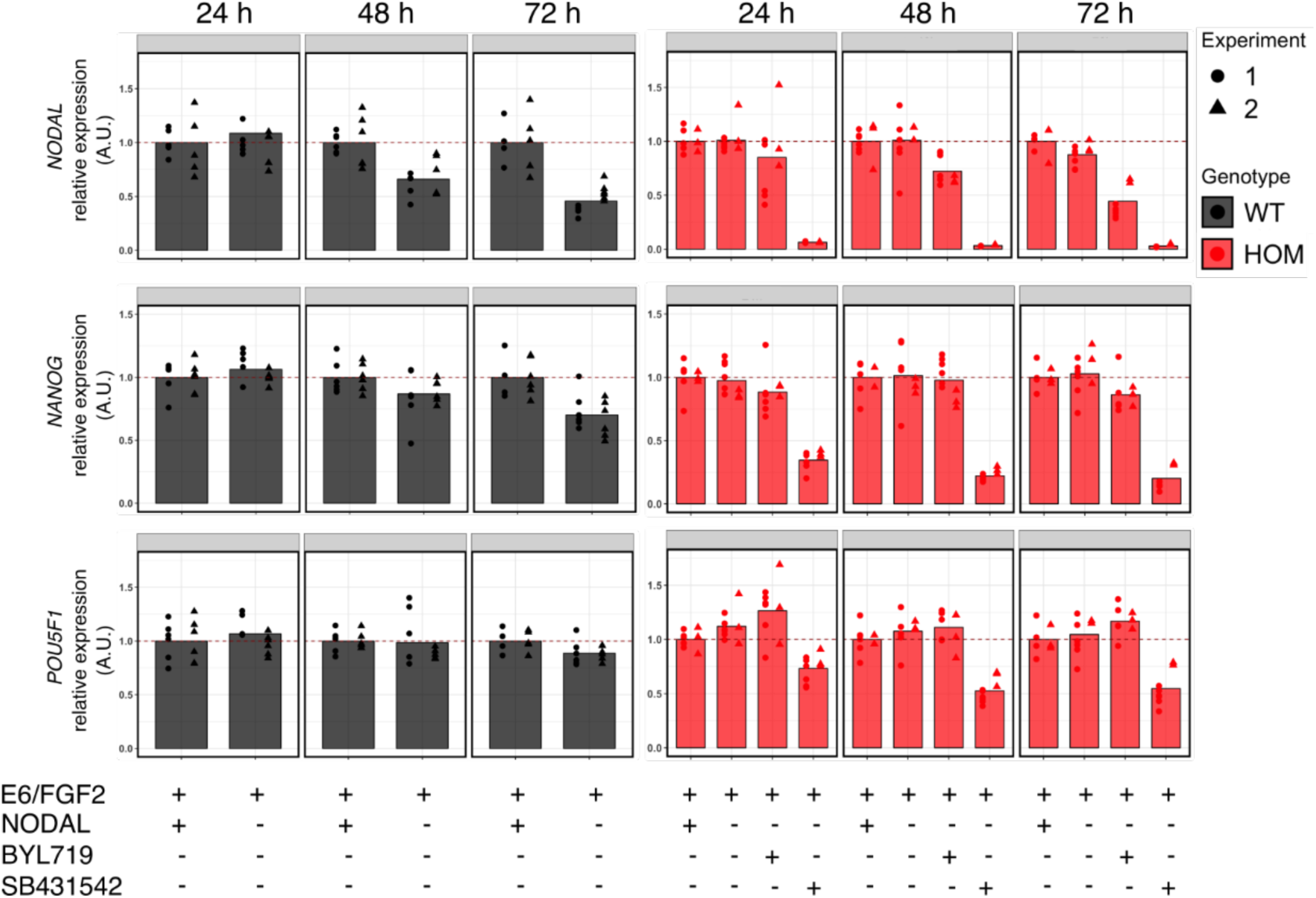
TGFβ signalling-dependent regulation of stemness in *PIK3CA*^*H1047R/H1047R*^ iPSCs. Gene expression time course of *NODAL, NANOG* and *POU5F1* in wild-type (WT) or *PIK3CA*^*H1047R/H1047R*^ iPSCs following the indicated treatments for 24 h, 48 h or 72 h. B250: 250 nM BYL719 (PI3Kα-selective inhibitor); E6/FGF2: Essential 6 medium supplemented with 10 ng/ml basic fibroblast growth factor 2 (FGF2). SB431542 is a specific inhibitor of the NODAL type I receptors ALK4/7 and the TGFβ type I receptor ALK5; used at 5 μM. When indicated, cultures were supplemented with 100 ng/ml NODAL. The data are from two independent experiments, with each treatment applied to triplicate cultures of three wild-type and two homozygous iPSC clones. To aid interpretation, gene expression values are normalised to the E6/FGF2 condition within each genotype and time-point. An alternative visualisation that illustrates the differential expression of *NODAL* and *NANOG* between mutant and wild-type cells is shown in Fig. S4A. For analysis of additional lineage markers, see Fig. S4B. For representative micrographs of *PIK3CA*^*H1047R/H1047R*^ iPSCs exposed to the different treatments, see Fig. S5. A.U., arbitrary units.

To confirm that TGFβ signalling is required for maintenance of stemness in *PIK3CA*^*H1047R/H1047R*^ iPSCs, the cells were treated with SB431542 – a specific inhibitor of TGFβ and NODAL type I receptors (34). This completely repressed *NODAL* expression within 24 h, accompanied by downregulation of *NANOG* expression (**Fig. 5**). A similar effect was observed on *POU5F1* expression, consistent with destabilisation of the pluripotency gene regulatory network in *PIK3CA*^*H1047R/H1047R*^ iPSCs (**Fig. 5**). Confirming this, we used a lineage-specific gene expression array to demonstrate similar reduction in expression of several other well-established stemness markers (*MYC, FGF4, GDF3*) with increased expression in *PIK3CA*^*H1047R/H1047R*^ iPSCs, performing the analysis after 48 h of TGFβ pathway inhibition (**Fig. S4B**). Despite the short treatment, we found evidence for the expected neuroectoderm induction upon inhibition of the TGFβ pathway (21, 35), reflected by increased expression of *CDH9, MAP2, OLFM3* and *PAPLN* (**Fig. S4B**).

Collectively, these data suggest that the stemness phenotype of *PIK3CA*^*H1047R/H1047R*^ iPSCs is mediated by self-sustained TGFβ signalling, most likely through PI3K dose-dependent increase in *NODAL* expression, and that this is amenable to reversal through inhibition of the TGFβ pathway but not of PI3Kα itself.

## Discussion

*PIK3CA*^*H1047R*^ is the most common activating *PIK3CA* mutation in human cancers and in PROS (6). We recently found that *PIK3CA*-associated cancers often harbour multiple mutated *PIK3CA* copies, and demonstrated that homozygosity but not heterozygosity for *PIK3CA*^*H1047R*^ leads to self-sustained stemness in human pluripotent stem cells (hPSCs) (7). High-depth transcriptomics in this study confirmed that heterozygosity for *PIK3CA*^*H1047R*^ induces significant but very modest transcriptional changes; observed both in CRISPR-edited hPSCs with long-term *PIK3CA*^*H1047R*^ expression and in mouse embryonic fibroblasts (MEFs) upon acute *PIK3CA*^*H1047R*^ induction by *Cre*, with canonical PI3K pathway activation seen in both cases (current study and Ref. (7, 26, 36)). Similarly, hPSCs with heterozygous expression of *PIK3CA*^*E418K*^, a “non-hotspot” mutation, were transcriptionally indistinguishable from their isogenic wild-type controls. In contrast to the mild transcriptional consequences of these heterozygous variants, however, homozygosity for *PIK3CA*^*H1047R*^ was associated with differential expression of nearly a third of the hPSC transcriptome, suggesting widespread epigenetic reprogramming. This near-binary response is not a consequence of a similar quantitative difference in PI3K pathway activation, as assessed by phosphoprotein profiling, which instead showed a relatively modest and graded increase in homozygous *versus* heterozygous *PIK3CA*^*H1047R*^ hPSCs. This implies that the apparent sharp PI3K signalling threshold that determines the cellular response in hPSCs is “decoded” distal to the canonical pathway activation.

Using a combination of computational analyses and targeted experiments, the current study further provides evidence for self-sustained TGFβ pathway activation as the main mechanism through which *PIK3CA*^*H1047R*^ homozygosity “locks” hPSCs in a differentiation-resistant state that has become independent of the driver mutation and the associated PI3K pathway activation. We suggest that homozygosity but not heterozygosity for *PIK3CA*^*H1047R*^ promotes sufficient TGFβ pathway activity to induce increased *NODAL* and downstream *NANOG* expression to levels that stabilise the stem cell state, yet are not high enough to tip the balance towards mesendoderm differentiation (7). Exactly how PI3K activation regulates *NODAL* expression remains unknown. A potential mechanism involves increased expression of the stem cell reprogramming factor MYC, which was observed at both mRNA and protein level in homozygous but not heterozygous *PIK3CA*^*H1047R*^ iPSCs. Furthermore, MYC was the only node in the WGCNA-based network of pluripotency, PI3K and TGFβ pathway components that was shared by all three pathways (**Fig. 4D**). MYC has previously been shown to exert oncogenic effects that depend on a sharp threshold of MYC expression, reminiscent of the effects we observe for dose-dependent *PIK3CA* activation (37). Elevated MYC has also been shown to allow *PIK3CA*^*H1047R*^-induced murine breast cancers to become independent of continuous *PIK3CA*^*H1047R*^ expression (38).

Stabilisation of the stemness phenotype in hPSCs by strong genetic PI3K pathway activation may be generalisable beyond the iPSC model system. BYL719 (alpelisib; Novartis), the PI3Kα-selective inhibitor used in our cellular studies, was recently approved for use in combination with anti-estrogen therapy in ER-positive breast cancers (39). In a separate study focusing on human breast cancer (Madsen *et al.*, manuscript submitted), we have described use of computational analyses to demonstrate a strong, positive relationship between a transcriptomically-derived PI3K activity score, stemness gene expression and tumour grade in breast cancer. Prior reports have suggested a role for NODAL in driving breast cancer stemness and aggressive disease (40, 41), with potential links to mTORC1 activation (42, 43). Our findings that BYL719 fails to fully reverse the increased *NODAL* and stemness gene expression in homozygous *PIK3CA*^*H1047R*^ iPSCs suggests that inhibition of TGFβ signalling as a pro-differentiation therapy warrants investigation as co-therapy with PI3K inhibitors in breast tumours with strong PI3K pathway activation. Finally, the lack of widespread transcriptional changes upon heterozygous expression of mutant *PIK3CA* in otherwise genetically normal cell models may explain the low oncogenicity of this genotype in isolation *in vivo*.

## Materials and Methods

All cell lines used in this study are listed in **Table S1**. Unless stated otherwise, standard chemicals were acquired from Sigma Aldrich, with details for the remaining reagents included in **Table S2**.

### iPSC culture and treatments

#### Maintenance

The derivation of the iPSC lines, including associated ethics statements, has been described previously (7). All lines were grown at 37°C and 5% CO2 in Essential 8 Flex (E8/F) medium on Geltrex-coated plates, in the absence of antibiotics. For maintenance, cells at 70-90% confluency were passaged as aggregates with ReLeSR, using E8 supplemented with RevitaCell (E8/F+R) during the first 24 h to promote survival. A detailed version of this protocol is available via protocols.io (doi: dx.doi.org/10.17504/protocols.io.4rtgv6n).

All cell lines were tested negative for mycoplasma and genotyped routinely to rule out cross-contamination during prolonged culture. Short tandem repeat profiling was not performed. All experiments were performed on cells within 10 passages since thawing.

#### Collection for RNA sequencing and total proteomics

For RNA sequencing and total proteomics, subconfluent cells were fed fresh E8/F 3 h prior to snap-freezing on dry ice and subsequent RNA or protein extraction. Relative to the results in Ref. (7), the current transcriptomic data of *PIK3CA*^*H1047R*^ were obtained more than 6 months following the first study, with cells at different passages, and were thus independent from one another. Moreover, sample collection for the second transcriptomics experiment was conducted over three days according to a block design, thus allowing us to determine transcriptional differences that are robust to biological variability.

#### Cell lysate collection for RPPA

For RPPA in growth factor-replete conditions, cells were fed fresh E8/F 3h before collection. To assess variability due to differences in collection timing, clones from each iPSC genotype were collected on each one of three days according to a block design, giving rise to a total of 22 cultures. To test the effect of the PI3Kα-specific inhibitor BYL719, cells were treated with 100 nM drug (or DMSO only as control treatment) for 24 h and exposed to growth factor removal within the last hour before collection. All cells were washed in DPBS prior to collection to rinse off residual proteins and cell debris.

#### TGFβ/NODAL signalling studies

Wild-type or homozygous *PIK3CA*^*H1047R*^ iPSCs were seeded in 12-well plates all coated with Geltrex from the same lot (#2052962; diluted in DMEM/F12 lot #RNBH0692). Cells were processed for seeding at a ratio of 1:15 according to the standard maintenance protocol. One day after seeding, individual treatments were applied to triplicate wells. Briefly, cells were first washed twice with 2 and 1 ml of Dulbecco’s PBS (DPBS) to remove residual growth factors. The base medium for individual treatments was Essential 6 supplemented with 10 ng/ml heat-stable FGF2. This was combined with one of the following reagents or their diluent equivalents: 100 ng/ml NODAL (diluent: 4 mM HCl), 250 nM BYL719 (diluent: DMSO), 5 μM SB431542 (diluent: DMSO). Cells were snap-frozen on dry ice after 24, 48 and 72 h following a single DPBS wash. Individual treatments were replenished daily at the same time of day to limit temporal confounders.

### Mouse embryonic fibroblast (MEF) culture

The derivation and culture of the wild-type and *PIK3CA*^*WT/H1047R*^ MEFs used in this study have been reported previously (26). Cell pellets were collected on dry ice 48 h after induction of heterozygous *PIK3CA*^*H1047R*^ expression, without prior starvation.

### RNA sequencing

Induced pluripotent stem cell lysates were collected in QIAzol and processed for RNA extraction with the DirectZol Kit as per the manufacturer’s instructions. The final RNA was subjected to quantification and quality assessment on an Agilent Bioanalyzer using the RNA 6000 Nano Kit, confirming that all samples had a RIN score of 10. For *PIK3CA*^*H1047R*^ iPSCs and corresponding wild-types, an Illumina TruSeq Stranded mRNA Library Prep Kit was used to synthesise 150-bp-long paired-end mRNA libraries, followed by sequencing on an Illumina HiSeq 4000, with average depth of 70 million reads per sample. *PIK3CA*^*WT/E418K*^ and isogenic control iPSCs were subjected to 50-bp-long single-end RNA sequencing (RNAseq) at an average depth of 20 million reads per sample.

MEF RNA was extracted using Qiagen’s RNAeasy miniprep (with QIAshredder). All samples had a confirmed Agilent Bioanalyzer RIN score of 10. An Illumina TruSeq Unstranded mRNA kit was used to prepare 100-bp-long paired-end libraries, followed by Illumina HiSeq 2000 sequencing.

Details of the subsequent data analyses (raw read mapping, counting, statistical testing, pathway and network analyses) are provided in SI Appendix.

### Label-free total proteomics

#### Sample preparation

Cells were cultured to subconfluence in Geltrex-coated T175 flasks, and protein was harvested by lysis in 3 ml modified RIPA buffer (50 mM Tris-HCl pH 7.5, 150 mM NaCl, 1% NP-40, 0.5% Na-deoxycholate, 1 mM EDTA) supplemented with phosphatase inhibitors (5 mM ß-glycerophosphate, 5 mM NaF, 1 mM Na_3_VO_4_) and protease inhibitors (Roche cOmplete ULTRA Tablets, EDTA-free). The lysates were sonicated on ice (4x 10s bursts, amplitude = 60%; Bandelin Sonopuls HD2070 sonicator) and spun down for 20 min at 4300g. Ice-cold acetone was added to the supernatant to achieve a final concentration of 80% acetone, and protein was left to precipitate overnight at -20°C. Precipitated protein was pelleted by centrifugation at 2000g for 5 min and solubilised in 6 M urea, 2 M thiourea, 10 mM HEPES pH 8.0. Protein was quantified using the Bradford assay and 8 mg of each sample were reduced with 1 mM dithiothritol, alkylated with 5 mM chloroacetamide and digested with endopeptidase Lys-C (1:200 v/v) for 3 h. Samples were diluted to 1 mg/ml protein using 50 mM ammonium bicarbonate and incubated overnight with trypsin (1:200 v/v). Digested samples were acidified and urea removed using SepPak C18 cartridges. Peptides were eluted, and an aliquot of 100 μg set aside for total proteome analysis. The peptides were quantified using the Pierce quantitative colorimetric peptide assay. The equalised peptide amounts were lyophilised and resolubilised in 2% acetonitrile and 1% trifluoroacetic acid in order to achieve a final 2 μg on-column peptide load.

#### Mass spectrometry (MS) data acquisition

All spectra were acquired on an Orbitrap Fusion Tribrid mass spectrometer (Thermo Fisher Scientific) operated in data-dependent mode coupled to an EASY-nLC 1200 liquid chromatography pump (Thermo Fisher Scientific) and separated on a 50 cm reversed phase column (Thermo Fisher Scientific, PepMap RSLC C18, 2 μM, 100A, 75 μm x 50 cm). Proteome samples (non-enriched) were eluted over a linear gradient ranging from 0-11% acetonitrile over 70 min, 11-20% acetonitrile for 80 min, 21-30% acetonitrile for 50 min, 31-48% acetonitrile for 30 min, followed by 76% acetonitrile for the final 10 min with a flow rate of 250 nl/min.

Survey-full scan MS spectra were acquired in the Orbitrap at a resolution of 120,000 from m/z 350-2000, automated gain control (AGC) target of 4×10^5^ ions, and maximum injection time of 20 ms. Precursors were filtered based on charge state (≥2) and monoisotopic peak assignment, and dynamic exclusion was applied for 45s. A decision tree method allowed fragmentation for ion trap MS2 via electron transfer dissociation (ETD) or higher-energy collision dissociation (HCD), depending on charge state and m/z. Precursor ions were isolated with the quadrupole set to an isolation width of 1.6 m/z. MS2 spectra fragmented by ETD and HCD (35% collision energy) were acquired in the ion trap with an AGC target of 1e4. Maximum injection time for HCD and ETD was 80 ms for proteome samples.

Details of the subsequent data analyses (FASTA file generation, mass spectrometry searches) are provided in SI Appendix.

### Reverse phase protein array (RPPA)

For RPPA, snap-frozen cells were lysed in ice-cold protein lysis buffer containing: 50 mM HEPES, 150 mM NaCl, 1.5 mM MgCl_2_, 10% (v/v) glycerol, 1% (v/v) TritonX-100, 1 mM EGTA, 100 mM NaF, 10 mM Na_4_P_2_O_7_, 2 mM Na_3_VO_4_ (added fresh), 1X EDTA-free protease inhibitor tablet, 1X PhosStop tablet. Protein concentrations were measured using BioRad’s DC protein assay, and all concentrations were adjusted to 1 mg/ml with lysis buffer and 1X SDS sample buffer (10% glycerol, 2% SDS, 62.5 mM Tris-HCl pH 6.8) supplemented with 2.5% β-mercaptoethanol.

The protein lysates were processed for slide spotting and antibody incubations as described previously (44). Briefly, a four-point dilution series was prepared for each sample and printed in triplicate on single pad Avid nitrocellulose slides (Grace Biolabs) consisting of 8 arrays with 36×12 spots each. Next, slides were blocked and incubated in primary and secondary antibodies. The processed arrays were imaged using an Innopsys 710 slide scanner. Non-specific signals were determined for each slide by omitting primary antibody incubation step. For normalisation, sample loading on each array was determined by staining with Fast Green dye and recording the corresponding signal at 800 nm. Details for all primary and secondary RPPA antibodies are included in **Table S3**.

Details of all subsequent data analyses, including statistical testing, are provided in SI Appendix.

### Reverse transcription-quantitative PCR (RT-qPCR)

Cellular RNA was extracted as described above for RNA Sequencing, and 200 ng used for complementary DNA (cDNA) synthesis with Thermo Fisher’s High-Capacity cDNA Reverse Transcription Kit. Subsequent SYBR Green-based qPCRs were performed on 2.5 ng total cDNA. TaqMan hPSC Scorecards (384-well) were used according to the manufacturer’s instructions with minor modifications. Further details on protocol modifications and all data analysis steps are provided in SI Appendix.

### Statistical analyses

Bespoke statistical analyses are specified in the relevant sections above and in SI Appendix.

### Data and materials availability

Raw data and bespoke RNotebooks containing guided scripts and plots are available *via* the Open Science Framework (doi: 10.17605/OSF.IO/MUERY). The original RNAseq data have been deposited to the Gene Expression Omnibus (GEO), under accession numbers: GSE134076 (H1047R iPSC data), GSE138161 (E418K iPSC data), GSE135046 (MEF data). The mass spectrometry proteomics data have been deposited to the ProteomeXchange Consortium via the PRIDE (45) partner repository with the dataset identifier PXD014719 (password to be provided to reviewers before public release). Further information and requests for resources and reagents should be directed to and will be fulfilled by the corresponding authors, Ralitsa R. Madsen or Robert K. Semple.

## Acknowledgements

We thank Dominique McCormick and Ineke Luijten for help with RNA extraction and cDNA synthesis, Cornelia Gewert for technical support, and Marcella Ma, Brian Lam and Michelle Dietzen for technical support with RNA sequencing and genomics analyses, respectively. We thank Evelyn K. Lau for help with GEO upload of the MEF RNAseq data. We are grateful to Prof Siddhartan Chandran and his group for iPSC culturing facilities and to Pamela Brown (SURF Biomolecular Core, University of Edinburgh) for access to qPCR facilities.

