## Supplementary Material for "NODAL/TGFβ signalling mediates the self-sustained stemness induced by *PIK3CA*^*H1047R*^ homozygosity in pluripotent stem cells"

**This PDF file includes:**

Supplementary text
Figures S1 to S5
Tables S1 to S4
Legends for Datasets S1 to S8
List of OSF-deposited source data and code
References

**Other supplementary materials for this manuscript include the following:**

Datasets S1 to S8
OSF-deposited source data and code (doi: 10.17605/OSF.IO/MUERY)

### Supplementary Material Text

#### iPSC processing for individual experiments

*Cell lysate collection for RNA sequencing and total proteomics*

For RNA sequencing and total proteomics, subconfluent cells were fed fresh E8/F 3 h prior to snap-freezing on dry ice and subsequent RNA or protein extraction. Relative to the results in Ref. (1), the current transcriptomic data of PIK3CAH1047R were obtained more than 6 months following the first study, with cells at different passages, and were thus independent from one another. Moreover, sample collection for the second transcriptomics experiment was conducted over three days according to a block design, thus allowing us to determine transcriptional differences that are robust to biological variability.

*Cell lysate collection for RPPA*

For RPPA in growth factor-replete conditions, cells were fed fresh E8/F 3h before collection. To assess variability due to differences in collection timing, clones from each iPSC genotype were collected on each one of three days according to a block design, giving rise to a total of 22 cultures. To test the effect of the PI3Kα-specific inhibitor BYL719, cells were treated with 100 nM drug (or DMSO only as control treatment) for 24 h and exposed to growth factor removal within the last hour before collection. All cells were washed in DPBS prior to collection to rinse off residual proteins and cell debris.

*TGFβ/NODAL signalling studies*

Wild-type or homozygous PIK3CAH1047R iPSCs were seeded in 12-well plates all coated with Geltrex from the same lot (#2052962; diluted in DMEM/F12 lot #RNBH0692). Cells were processed for seeding at a ratio of 1:15 according to the standard maintenance protocol. One day after seeding, individual treatments were applied to triplicate wells. Briefly, cells were first washed twice with 2 and 1 ml of Dulbecco's PBS (DPBS) to remove residual growth factors. The base medium for individual treatments was Essential 6 supplemented with 10 ng/ml heat-stable FGF2. This was combined with one of the following reagents or their diluent equivalents: 100 ng/ml NODAL (diluent: 4 mM HCl), 250 nM BYL719 (diluent: DMSO), 5 μM SB431542 (diluent: DMSO). Cells were snap-frozen on dry ice after 24, 48 and 72 h following a single DPBS wash. Individual treatments were replenished daily at the same time of day to limit temporal confounders.

#### RNA sequencing data analyses

*Raw read mapping, counting and differential expression*

Raw reads were mapped to the human genome build GRCh38 (for iPSC RNAseq) or the mouse genome build GRCm38 (for MEF RNAseq), and gene level counts were performed using Spliced Transcripts Alignment to a Reference (STAR) v2.5 (2). Subsequent data processing was performed using open-source R software according to the *limma-voom* method (3). Briefly, raw counts were converted to counts per million (cpm) using the *cpm()* function in the *edgeR* package (4), followed by normalisation according the trimmed mean of M (TMM) method (5). The mean-variance relationship was modelled with *voom()*, followed by linear modelling and computation of moderated t-statistics using the *lmFit()* and *eBayes()* functions in the *limma* package (3). The associated P-value for assessment of differential gene expression was adjusted for multiple comparisons with the Benjamini-Hochberg method at false-discovery rate (FDR) $\leq$ 5% (6). The function *duplicateCorrelation()* was applied to correct for the use of replicate iPSC clones.

Correlations between corresponding transcriptomics and/or proteomics data were calculated using Spearman's rank-order correlation test for non-normally distributed data.

#### Pathway and network analyses

*Ingenuity® Pathway Analysis (IPA)*

The list of differentially expressed total proteins (*PIK3CA*$^{H1047R/H1047R}$ *vs* wild-type) was subjected to IPA (build version: 448560M; content version: 36601845) against the Ingenuity Knowledge Base, considering only relationships where confidence was classified as "Experimentally Observed". Following exclusion of chemicals and drugs, the Upstream Regulators list was used for generation of Volcano plots of the respective activation z-scores and overlap p-values.

IPA was also used to analyse the lists of differentially expressed genes in both heterozygous and homozygous *PIK3CA*$^{H1047R}$ iPSCs (IPA build version: 484108M; content version: 45868156) and MEFs (IPA build version: 486617M; content version: 46901286), using the Ingenuity Knowledge Base and considering only relationships where confidence was classified as "Experimentally Observed". Chemicals and diseases were excluded from Node Types. For the iPSC datasets, differentially expressed genes were only considered for IPA analysis if having an absolute log2(fold-change) $\geq$ log2(1.3). The choice of this relatively permissive log fold-change choice was guided by the *limma()* tutorial for RNAseq (3), and the assumption that small fold-changes in the expression of genes that act within the same pathway may be sufficient to elicit a functionally important response and thus should not be omitted. This consideration is of particular relevance for transcriptomic data from heterozygous *PIK3CA*$^{H1047R}$ iPSCs where fold-changes were relatively small. Due to the high number of differentially expressed transcripts in *PIK3CA*$^{H1047R/H1047R}$ iPSCs, the analysis was conducted using the top 2000 up- and top 2000 downregulated transcripts.

The IPA Upstream Regulator Analysis is based on the proprietary Ingenuity Knowledge Base which is used to compute two scores based on user-specified data: an enrichment score (Fisher's exact test p-value) that measures overlap between observed and predicted regulated gene sets; a z-score that assesses the match between observed and predicted up/down regulation patterns (7). The results of the Upstream Regulators Analysis were extracted for downstream Volcano plotting of overlap p-values and associated activation z-scores. Note that for heterozygous *PIK3CA*$^{H1047R}$ iPSCs, a bias-corrected activation z-score was used for plotting to take into account any bias arising from a larger number of upregulated *vs* downregulated genes in these cells.

*Weighted Gene Correlation Network Analysis (WGCNA)*

RNA sequencing counts from all 12 samples were converted to reads per kilobase million (RPKM). A threshold of 10 RPKM was used to filter out low-expression genes, followed by removing any genes with missing values caused by this filtering. The RPKM values for the remaining 16,823 genes were log2 transformed and taken forward for network analysis using the *WGCNA* R package (8, 9), with a soft power threshold of 28 (chosen to maximise scale independence and minimise mean connectivity) and a minimum module size of 30 genes.

To identify modules associated with homozygosity for *PIK3CA*$^{H1047R}$, we used the correlation between a gene's module membership (eigengene) and significance for differential expression in homozygous *PIK3CA*$^{H1047R/H1047R}$ iPSCs. The top two most significant modules for the homozygosity trait were selected for functional enrichment analysis. CytoScape plugin ClueGO (version 2.5.4) (10) was used to perform pathway analysis using the Kyoto Encyclopedia of Genes and Genomes (KEGG) ontology (build 27.02.19) (11). All settings were kept at default values. Only pathways with p-value $\leq$ 0.05 were selected, and a custom reference gene set was used as background (the 16,823 genes analysed using WGCNA). Network visualisation was performed using Cytoscape (12).

**Mass spectrometry data acquisition and analyses**

*Mass spectrometry (MS) data acquisition*

All spectra were acquired on an Orbitrap Fusion Tribrid mass spectrometer (Thermo Fisher Scientific) operated in data-dependent mode coupled to an EASY-nLC 1200 liquid chromatography pump (Thermo Fisher Scientific) and separated on a 50 cm reversed phase column (Thermo Fisher Scientific, PepMap RSLC C18, 2 $\mu$M, 100A, 75 $\mu$m x 50 cm). Proteome samples (non-enriched) were eluted over a linear gradient ranging from 0-11% acetonitrile over 70 min, 11-20% acetonitrile for 80 min, 21-30% acetonitrile for 50 min, 31-48% acetonitrile for 30 min, followed by 76% acetonitrile for the final 10 min with a flow rate of 250 nl/min.

Survey-full scan MS spectra were acquired in the Orbitrap at a resolution of 120,000 from m/z 350-2000, automated gain control (AGC) target of 4x105 ions, and maximum injection time of 20 ms. Precursors were filtered based on charge state (≥2) and monoisotopic peak assignment, and dynamic exclusion was applied for 45s. A decision tree method allowed fragmentation for ion trap MS2 via electron transfer dissociation (ETD) or higher-energy collision dissociation (HCD), depending on charge state and m/z. Precursor ions were isolated with the quadrupole set to an isolation width of 1.6 m/z. MS2 spectra fragmented by ETD and HCD (35% collision energy) were acquired in the ion trap with an AGC target of 1e4. Maximum injection time for HCD and ETD was 80 ms for proteome samples.

*Whole-exome sequencing (WES) and FASTA file generation*

WES was performed on a single clone per genotype to generate cell-specific databases for downstream mass spectrometry searchers. Genomic DNA was extracted with Qiagen's QIAamp DNA Micro Kit according to the manufacturer's instructions, followed by quantification using the Qubit dsDNA High Sensitivity Assay Kit and by dilution to 5 ng/µl in the supplied TE buffer. The samples were submitted for library preparation and sequencing by the SMCL Next Generation Sequencing Hub (Academic Laboratory of Medical Genetics, Cambridge). Sequencing was performed on an Illumina HiSeq 4000 with 50X coverage across more than 60% of the exome in each sample. Raw reads were filtered with Trimmomatic (13) using the following parameters: headcrop = 3, minlen = 30, trailing = 3. The trimmed reads were aligned to the human reference genome (hg19 build) with BWA (14), followed by application of GATK base quality score recalibration, indel realignment, duplicate removal and SNP/indel discovery with genotyping (15). GATK Best Practices standard hard filtering parameters were used throughout (16).

In order to find non-reference, mutated peptides in the MS data, we increased the search FASTA file with mutations affecting the protein sequence, as detected by WES with a high sensitivity filter: QD $\leq$ 1.5, FS $\geq$ 60, MQ $\geq$ 40, MQRankSum $\leq$ -12.5, ReadPosRankSum $\leq$ -8.0, and average DP $\geq$ 5 per sample. The Ensembl Variant Effect Predictor (VEP) with Ensembl v88 was used to predict the effect of the mutations on the protein sequence (17). For every variant with an effect on the protein sequence we added the predicted mutated tryptic peptide at the end of the protein sequence.

*Mass spectrometry searches*

Raw files were processed using MaxQuant 1.5.0.2 (18) with all searches conducted using cell-specific databases (see *Whole-exome sequencing and FASTA file generation*), where all protein sequence variants were included in addition to the reference (Ensemble v68 human FASTA). Methionine oxidation, protein N-terminal acetylation and serine/threonine/tyrosine phosphorylation were set as variable modifications and cysteine carbamidomethylation was set as a fixed modification. False discovery rates were set to 1% and the "match between runs" functionality was activated. We filtered out peptides that were associated with multiple identifications in the MaxQuant *msms.txt* file, had a score < 40, were identified in the reverse database or came from known contaminants. Analysis of the observed peptides passing these filters was performed using a Monte Carlo Markov Chain model as described previously (19). Briefly, the model predicted the average ratio (sample versus control) of a peptide as a function of the observed protein concentration (obtained from the MaxQuant *evidence.txt* file). Combined with a noise model, a distribution of likely values for the parameters was obtained. The mean and standard deviation of this resulting distribution was used to calculate a z-score which was used together with the fold-change (FC) for subsequent filtering for differentially expressed proteins ($|z| \geq 1.2$;$|\ln(FC)| \geq \ln(1.2)$)

**RPPA data analyses**

Slide images were analysed using Mapix software (Innopsys), with the spot diameter of the grid set to 270 µm. Background signal intensity was determined for each spot individually and subtracted from the sample spot signal. A test for linearity was performed from the four-point dilution series, according to a flag system where $R^2 > 0.9$ was deemed good, $R^2 > 0.8$ was deemed acceptable and $R^2 < 0.8$ was poor (excluded from subsequent analyses). Median values from the four-point dilution series were calculated for each technical replicate and normalised to the corresponding Fast Green value to account for differences in protein loading. For each sample and protein target, a mean expression value was calculated from the remaining technical replicates and normalised to the corresponding mean of the wild-type group. All phosphoprotein signals were also normalised to the corresponding total protein values.

**RT-qPCR set-up and data analyses**

For SYBR Green-based qPCRs, A 5-fold cDNA dilution series was prepared and used as standard curve for relative quantitation of gene expression. *TBP* was used as normaliser following confirmation that its gene expression remained unaffected by the tested conditions. Melt curve analyses confirmed amplification of a single product by each primer pair. All primers had amplification efficiencies 95%-105%. Samples were loaded in duplicate in 384-well plates.

The TaqMan hPSC Scorecard was set up according to the manufacturer's instructions with the following modifications. From each cDNA sample diluted to 20 ng/μl, two 50 μl RT reactions were set up, with 500 ng RNA sample in each. Next, the two RT replicates were combined to obtain 1 μg cDNA in a total volume of 100 μl (final concentration: 10 ng/μl). This was subsequently diluted to 0.715 ng/μl and 10 μl loaded into each Scorecard well. All Ct values were mapped to their corresponding genes using the TaqMan hPSC Scorecard analysis software provided by the manufacturer. Genes with Ct values < 15 were excluded from further analyses. To be considered for downstream processing, genes were also required to have Ct values < 30 in at least two out of the eight samples. Next, Ct values were linearised (antilog) under the assumption of 100 % primer amplification efficiency. The geometric expression mean of the control gene assays was used for subsequent normalisation of individual gene expression values.

All qPCR data were acquired on a Quant Studio™ 5 Real-Time PCR System (Thermo Fisher Scientific). The thermocycling conditions (SYBR Green reactions) were as follows (ramp rate 1.6°C/s for all): 50°C for 2 min, 95°C for 10 min, 40 cycles at 95°C for 15 sec and 60°C for 1 min, followed by melt curve analysis (95°C for 15 sec, 60°C for 1 min, and 95°C for 15 min with ramp rate 0.075°C/sec). The TaqMan hPSC Scorecard thermocycling conditions were as specified by the manufacturer in the accompanying template.

All relevant primer sequences are included in **Table S4**.

**Statistical analyses**

In line with recent (ATOMIC) recommendations by the American Statistical Association (20), we have avoided arbitrary use of "statistical significance" applied to data from small-scale cell culture experiments which violate assumptions of the most widely used statistical tests. Instead, we present all data from multiple orthogonal experiments, alongside complete information on experimental replicates, independent clones and replicate cultures.

For RPPA data in Fig. 2A, a statistical test for differential expression was performed on datasets with more than three samples per group, using the *limma* package to apply the *limma-trend* method with *lmFit()* and *eBayes()*, specifying collection time as blocking factor (3). Phosphoprotein and total protein lists were processed separately. The associated p-value for assessment of differential gene expression was adjusted for multiple comparisons with the Benjamini-Hochberg method at FDR $\leq$ 5% (6). The function *duplicateCorrelation()* was applied to correct for the use of replicate iPSC clones on the same day. Heatmaps were generated using the *heatmap.2()* function within the *gplots* package in R, using target-wise correlation for dendrogram construction.

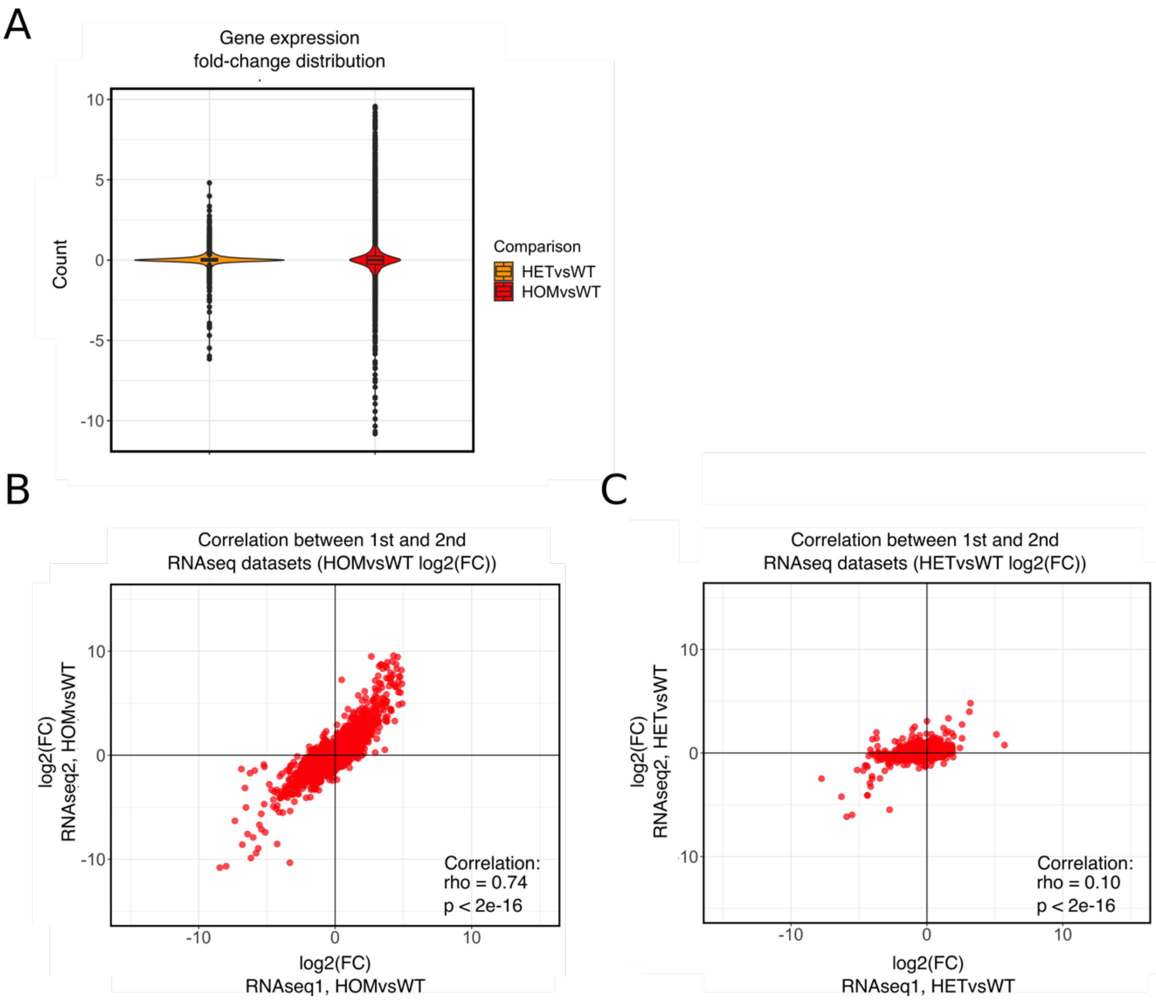


**Fig. S1, related to Fig. 1. Fold-change distribution and transcriptome correlations.** (A) Combined violin-boxplot representation of the fold-change (log2) distribution of gene expression changes between heterozygous and homozygous *PIK3CA*$^{H1047R}$ and wild-type iPSCs as indicated. (B) and (C) Correlation plot of the log2 expression fold-changes (FC) of significant total proteins *vs* the corresponding mRNA transcripts in *PIK3CA*$^{H1047R/H1047R}$ (HOM) (B) and *PIK3CA*$^{WT/H1047R}$ (HET) (C) iPSCs. Spearman's rho and the corresponding P-values are indicated on all plots. The first transcriptomic study (RNAseq1) was based on three independent cultures from three different clones per genotype; the second transcriptomic study (RNAseq2) used four independent cultures per genotype, from at least two independent clones each.

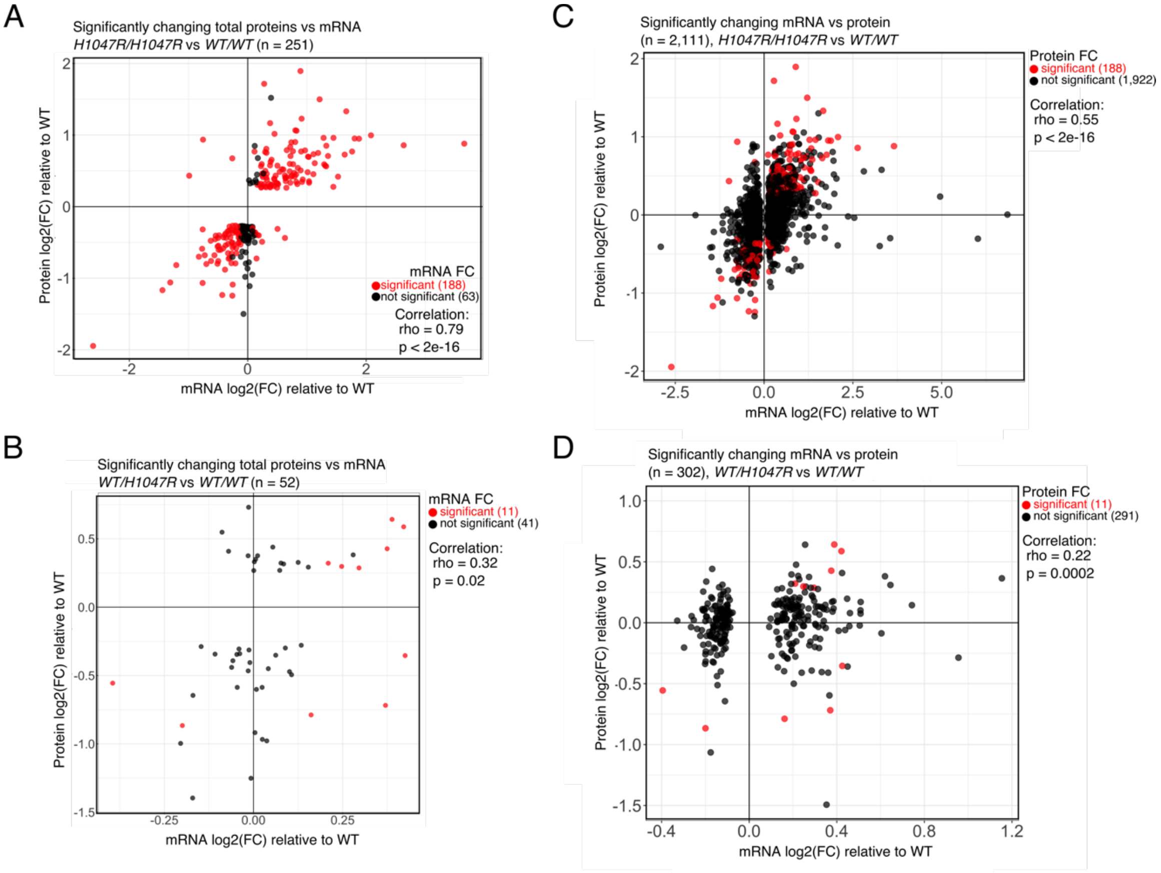


**Fig. S2**, **related to Fig. 1. Transcriptome-proteome correlations.** (A) Correlation plot of the log2 expression fold-changes (FC) of differentially expressed total proteins (|z| ≥ 1.2 and |ln(FC)| ≥ 1.2) in *PIK3CA*$^{H1047R/H1047R}$ iPSCs and the corresponding mRNA transcripts in an independent set of cultures. If identified as differentially expressed following statistical analysis (FDR ≤ 0.05), mRNA transcripts are highlighted in red irrespective of absolute fold-change. (B) As in (A) but starting with all differentially expressed mRNA transcripts (FDR ≤ 0.05 irrespective of fold-change magnitude) and plotting them to the corresponding protein identified by total proteomics. If differentially expressed (see (a)), the matched proteins are highlighted in red. (C) and (D) As in (A) and (B), respectively, but using the data for *PIK3CA*$^{WT/H1047R}$ iPSCs. All proteomic data were obtained from 3 independent clones per genotype using cultures at passages P47-P52, corresponding to the cultures used in our previous study (1). The high-depth transcriptomic data were obtained from 4 independent cultures (minimum 2 independent clones) per genotype using cultures at passages P55-P59. Spearman's rho and the corresponding P-values are indicated on all plots.

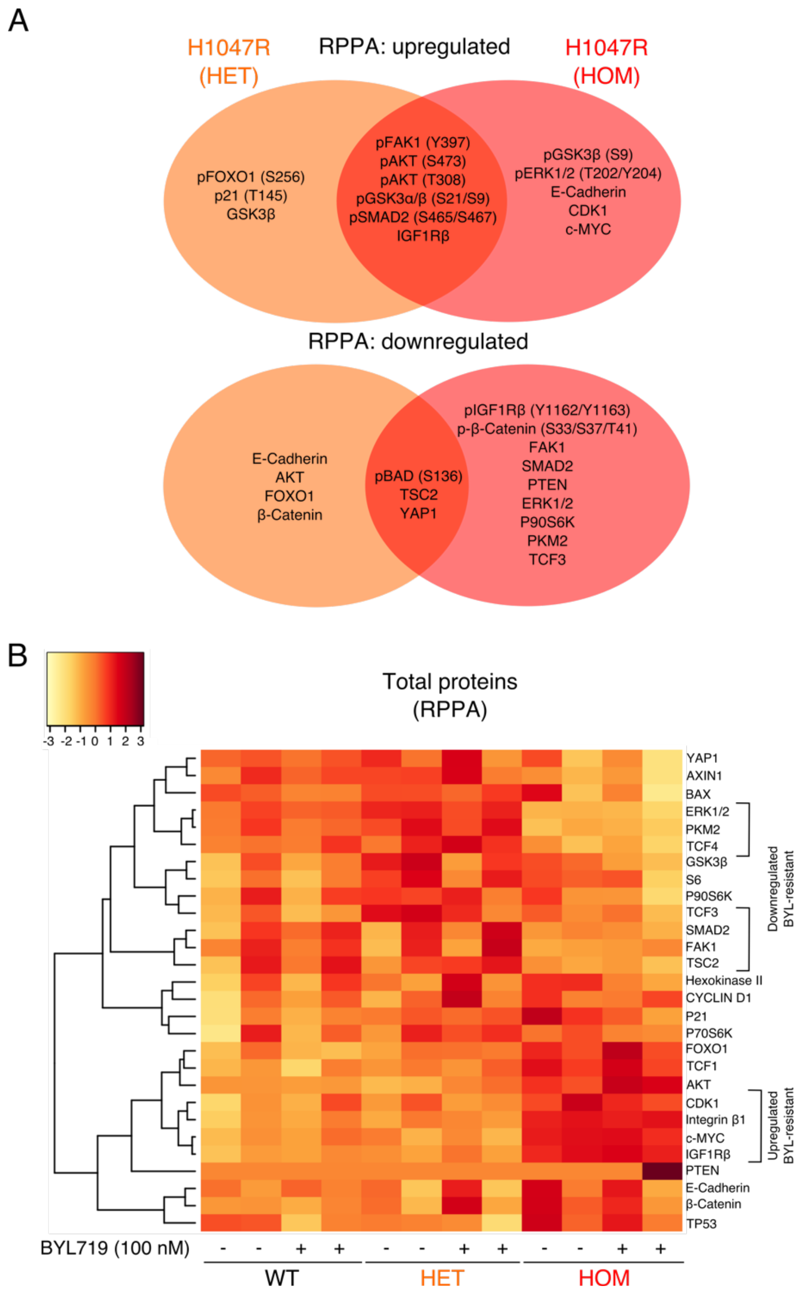


**Fig. S3, related to Fig. 2. Additional RPPA data.** (A) Venn-diagrams specifying differentially expressed phosphorylated and total proteins in *PIK3CA*$^{WT/H1047R}$ (HET) and *PIK3CA*$^{H1047R/H1047R}$ (HOM) iPSCs relative to wild-type controls, based on RPPA profiling of cells cultured in growth factor-replete conditions. The data are based on a total of 10 wild-type cultures, 5 *PIK3CA*$^{WT/H1047R}$ cultures and 7 *PIK3CA*$^{H1047R/H1047R}$ cultures, and all shown targets were differentially expressed at a false-discovery rate (FDR) $\leq$ 0.05. Following quality checks (see Materials and Methods), the RPPA data included 21 phosphorylated and 21 total proteins. (B) Heatmap of total proteins from RPPA profiling of wild-type (WT), *PIK3CA*$^{WT/H1047R}$ (HET) and *PIK3CA*$^{H1047R/H1047R}$ (HOM) iPSCs following short-term growth factor removal (1 h), +/- 100 nM BYL719 (PI3Kα inhibitor) for 24 h. Groups of total proteins exhibiting a consistent expression pattern in BYL719-treated *PIK3CA*$^{H1047R/H1047R}$ iPSCs are specified.

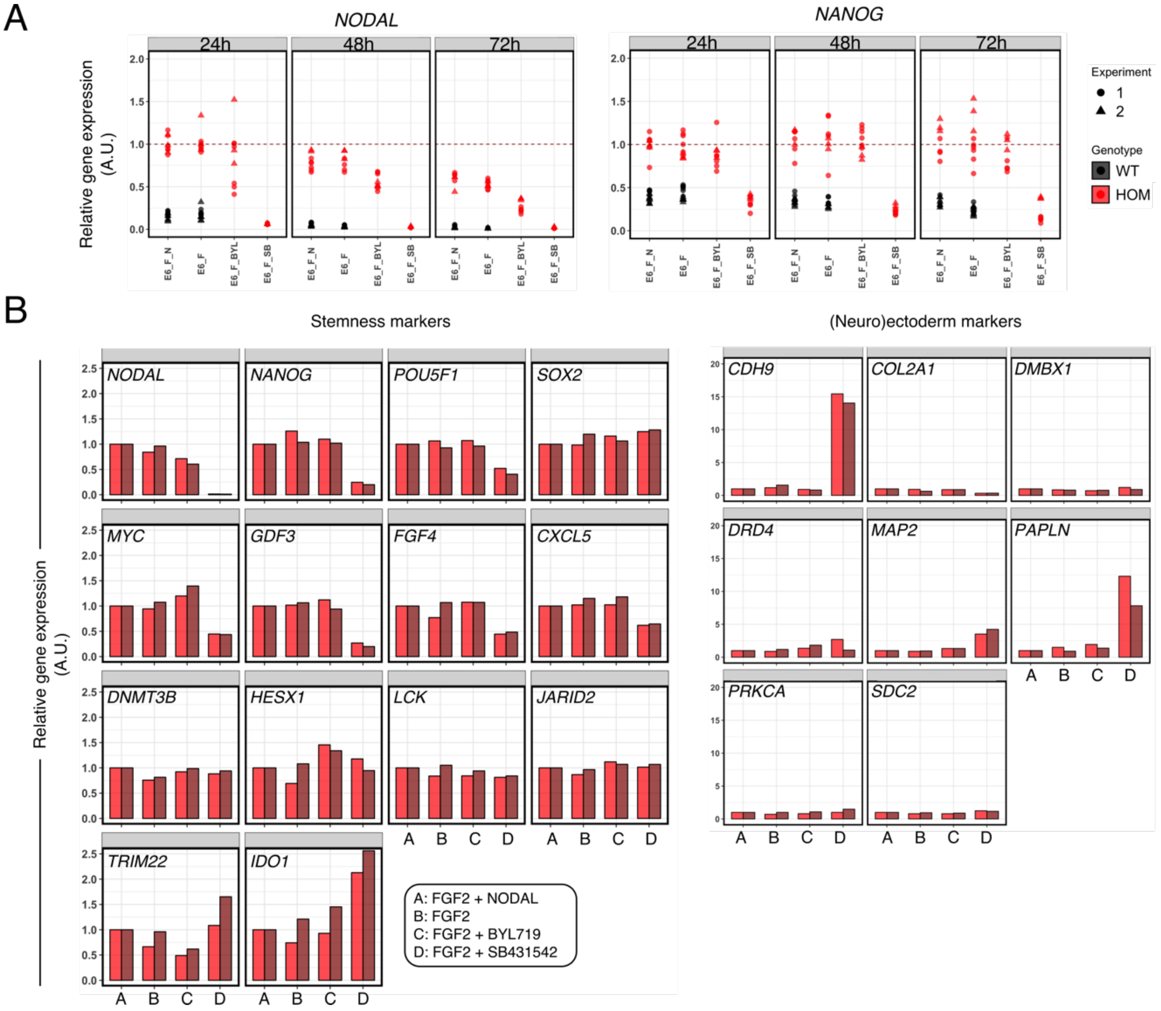


**Fig. S4**, **related to Fig. 5. Alternative representation of the experimental data in Fig. 5A and additional RT-qPCR-based profiling of lineage-specific markers.** (A) As in Fig. 5A but representing individual gene expression values for *NODAL* and *NANOG* scaled to the mean expression in *PIK3CA*$^{H1047R/H1047R}$ after 24 h in E6 medium supplemented with FGF2 and NODAL. A.U., arbitrary units. (B) TaqMan hPSC Scorecards were used to profile a set of stemness and (neuro)ectoderm markers in *PIK3CA*$^{H1047R/H1047R}$ iPSCs following the indicated treatments for 48 h. Each bar corresponds to a single sample, with colours specifying the use of two independent clones per treatment. For each gene and clone, values are scaled to the expression value in cells cultured in E6 medium with FGF2 and NODAL.

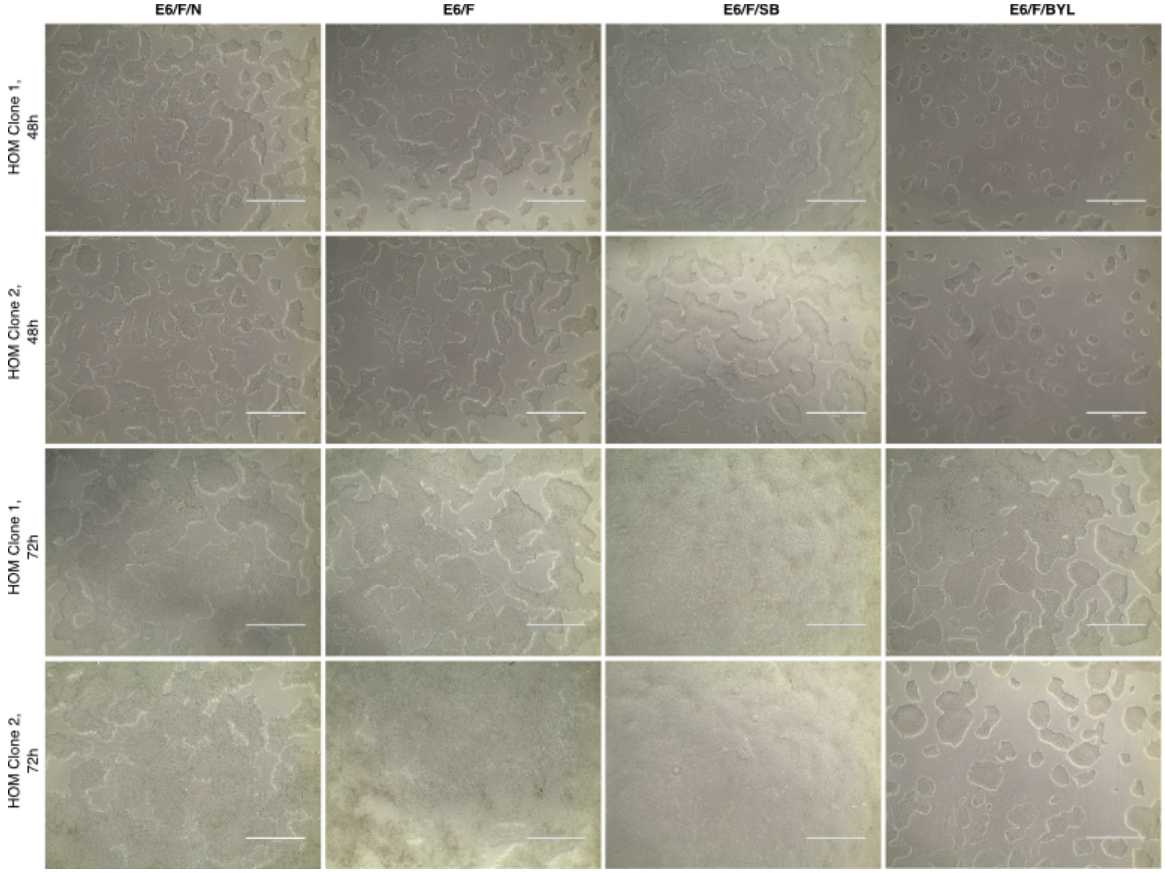


**Fig. S5**, **related to Fig. 5. Light micrographs of *PIK3CA*$^{H1047R/H1047R}$ iPSC clones exposed to different media formulations +/- specific inhibitors.** Each row corresponds to images of parallel cultures of one of two independent iPSC clones exposed to different treatments for 48 or 72 h as indicated. E6/F/N: Essential 6 medium supplemented with 10 ng/ml FGF2 and 100 ng/ml NODAL; E6/F: Essential 6 medium supplemented with 10 ng/ml FGF2; E6/F/SB: Essential 6 medium supplemented with 10 ng/ml FGF2 and 5 µM SB431542; E6/F/BYL: Essential 6 medium supplemented with 10 ng/ml FGF2 and 250 nM BYL719. The images are representative of 3 independent experiments. Scale bar: 1000 µM.

| Cell line | Source | Identifier |
|---|---|---|
| WTC11 male parental iPSC line | Coriell Institute | GM25256 |
| WTC11-derived male wild-type post-CRISPR editing | Madsen *et al.* PNAS 2019 (1) | N/A |
| WTC11-derived male *PIK3CA*$^{WT/H1047R}$ | Madsen *et al.* PNAS 2019 (1) | N/A |
| WTC11-derived male *PIK3CA*$^{H1047R/H1047R}$ | Madsen *et al.* PNAS 2019 (1) | N/A |
| M98-WT female iPSC | Madsen *et al.* PNAS 2019 (1) | N/A |
| M98-E418K (*PIK3CA*$^{WT/E418K}$) female iPSC | Madsen *et al.* PNAS 2019 (1) | N/A |
| Primary MEFs *PIK3CA*$^{WT/WT}$ (H1047R uninduced) | Moniz *et al.* Sci Reports 2017 (21) | N/A |
| Primary MEFs *PIK3CA*$^{WT/H1047R}$ (induced for 48h) | Moniz *et al.* Sci Reports 2017 (21) | N/A |

**Table S1. Cell lines.**

| Item | Catalogue no. | Vendor | Application |
|---|---|---|---|
| 100 ml DMSO, sterile-filtered | D2650 | Sigma Aldrich | Cells |
| 2X PowerUp SYBR Green Master Mix | A25742 | Thermo Fisher Scientific | Molbio |
| Acetonitrile | 1.00029 | Sigma Aldrich | Protein work |
| Amonium Bicarbonate | 09830 | Sigma Aldrich | Protein work |
| Bioanalyzer RNA 6000 Nano Kit | 5067-1511 | Agilent | Molbio |
| BYL719 (lot #2598155) | B9700-1mg | Cambridge Bioscience | Cells |
| Chloroacetamide | C0267 | Sigma Aldrich | Protein work |
| cOmplete ULTRA Tablets, EDTA-free | 5892791001 | Roche | Protein work |
| cOmplete ULTRA Tablets, Mini, EasyPack, PhosStop | 4906845001 | Roche | Protein work |
| DC Protein Assay Reagent A | 5000113 | Biorad | Protein |
| DC Protein Assay Reagent B | 5000114 | Biorad | Protein |
| DC Protein Assay Reagent S | 5000115 | Biorad | Protein |
| DirectZol RNA Miniprep Kit | R2051 | ZymoResearch | Molbio |
| Dithiothritol | 10197777001 | Sigma Aldrich | Protein work |
| DMEM/F12 without L-Glutamine, HEPES 500 ml | D6421 | Sigma Aldrich | Cells |
| DPBS without $Ca^{2+}$ and $Mg^{2+}$ | D8537 | Sigma Aldich | Cells |
| EndoFree Plasmid Maxi Kit (10) | 12362 | Qiagen | Molbio |
| Endopeptidase Lys-C | 129-02541 | Wako | Protein work |
| Essential 6 Medium | A1516401 | Thermo Fisher Scientific | Cells |
| Essential 8 Flex Medium Kit | A2858501 | Thermo Fisher Scientific | Cells |
| Geltrex LDEV Free hESC Quality 5 ml | A1413302 | Thermo Fisher Scientific | Cells |
| Gibco Heat Stable Recombinant Human bFGF (for prolonged culture), 100 ug (lot #2046190) | PHG0369 | Thermo Fisher Scientific | Cells |
| HEPES | H4034 | Sigma Aldrich | Protein work |
| High-Capacity cDNA Reverse Transcription Kit | 4368814 | Thermo Fisher Scientific | Molbio |
| InSolution™ TGF-b RI Kinase Inhibitor VI, SB431542 (lot #3075261) | 616464-5MG | Merck Millipore | Cells |
| Nuclease-free $H_2O$ | AM9932 | Thermo Fisher Scientific | Molbio/Portein work |
| Nunc™ Delta Cell-Culture Treated Multidishes; 6-well | 140685 | Thermo Fisher Scientific | Cells |
| Nunc™ Delta Cell-Culture Treated Multidishes; 12-well | 150628 | Thermo Fisher Scientific | Cells |
| Pierce Quantitative Colorimetric Peptide Assay | 23275 | Thermo Fisher Scientific | Protein work |

| | | | |
|---|---|---|---|
| QIAamp DNA Micro Kit (50) | 56304 | Qiagen | Molbio |
| QIAshredder | 79654 | Qiagen | Molbio |
| QIAzol Lysis Reagent | 79306 | Qiagen | Molbio |
| Quibit dsDNA HS Assay Kit | Q32854 | Thermo Fisher Scientific | Molbio |
| Recombinant human NODAL protein, 25 ug (lot #OLF1115091) | 3218-ND/CF | R&D | Cells |
| ReLeSR 100 ml | 5872 | Stem Cell Technologies | Cells |
| RevitaCell Suppplement 5 ml | A2644501 | Thermo Fisher Scientific | Cells |
| RNeasy Mini Kit | 74104 | Qiagen | Molbio |
| SepPak C18 cartridges | WAT020515 | Waters | Protein work |
| SuperScript IV Fist Strand Synthesis System | 18091050 | Thermo Fisher Scientific | Molbio |
| Urea | 15604 | Sigma Aldrich | Protein work |
| TaqMan® hPSC Scorecard™ Panel, 384-well | A15870 | Thermo Fisher Scientific | Molbio |
| Thiourea | T7875 | Sigma Aldrich | Protein work |
| Trifluoroacetic acid | T6508 | Sigma Aldrich | Protein work |
| TrueSeq Stranded mRNA Library Prep Kit | 20020594 | Illumina | Molbio |
| TruSeq RNA Library Prep Kit v2 | RS-930-2002 | Illumina | Molbio |
| TruSeq RNA Single Indexes Set A | 20020492 | Illumina | Molbio |
| Trypsin | T6567 | Sigma | Protein work |
| VenorGeM Classic Mycoplasma Kit | 11-1005 | Minerva Biolabs | Molbio |

**Table S2, Materials list**. Molbio, molecular biology.

| Antibody (clone if mAb) | Vendor | Catalogue # | Species |
|---|---|---|---|
| 4E-BP1 | CST | 9452 | rabbit |
| 4E-BP1 P Ser65 | CST | 9451 | rabbit |
| 4E-BP1 P Ser65 (174A9) | CST | 9456 | rabbit |
| 4E-BP1 P Thr37,Thr46 | CST | 9459 | rabbit |
| Acetyl-CoA carboxylase P Ser79 | CST | 3661 | rabbit |
| AKT | CST | 9272 | rabbit |
| AKT P Ser473 | CST | 9271 | rabbit |
| AKT P Thr308 | CST | 9275 | rabbit |
| AXIN1 (C76H11) | CST | 2087 | rabbit |
| β -Catenin | CST | 9562 | rabbit |
| β -Catenin P Ser33,Ser37,Thr41 | CST | 9561 | rabbit |
| β -Catenin P Thr41,Ser45 | CST | 9565 | rabbit |
| BAD | CST | 9292 | rabbit |
| BAD P Ser112 | CST | 9291 | rabbit |
| BAD P Ser136 | CST | 9295 | rabbit |
| BAX | CST | 2772 | rabbit |
| c-MYC | CST | 5605 | rabbit |
| c-MYC P Thr58,Ser62 | Epitomics | 1203-1 | rabbit |
| CDK1 | CST | 9112 | rabbit |
| CDK1 P Tyr15 | CST | 9111 | rabbit |
| Cyclin D1 | CST | 2926 | mouse |
| Cyclin D1 P Thr286 | CST | 3300 | rabbit |
| E-Cadherin | CST | 3195 | rabbit |
| ERK1/2 | CST | 9102 | rabbit |
| ERK1/2 P CST Thr202/Thr185,Tyr204/Tyr187 | CST | 4370 | rabbit |
| FAK1 | CST | 3285 | rabbit |
| FAK1 P Y397 | CST | 3283 | rabbit |
| FOXO1 P S256 | CST | 9461 | rabbit |
| FOXO1 (C29H4) | CST | 2880 | rabbit |
| GSK3α/β P Ser21/Ser9 | CST | 9331 | rabbit |
| GSK3β | CST | 9315 | rabbit |
| GSK3β P Ser9 | CST | 9336 | rabbit |
| Hexokinase II | CST | 2867 | rabbit |
| IGF1Rβ | CST | 3027 | rabbit |
| IGF1Rβ P Tyr1162,Tyr1163 | Invitrogen (Biosource) | 44-804G | rabbit |
| Integrin β 1 (EP1041Y) | abcam | ab52971 | rabbit |
| Lamin A/C | CST | 2032 | rabbit |
| LEF1 (C12A5) | CST | 2230 | rabbit |
| P21 | CST | 2946 | mouse |
| P21 p Thr145 | Santa Cruz | sc-20220-R | rabbit |
| p110α | CST | 4249 | rabbit |
| PKM2 XP(R) | CST | 4053 | rabbit |
| PKM2 P Tyr105 | CST | 3827 | rabbit |
| PTEN | CST | 9552 | rabbit |
| RB P Ser780 | CST | 9307 | rabbit |
| Rictor P T1135 | CST | 3806 | rabbit |
| RPS6KA (Rsk1-3) | Santa Cruz | sc-231 | rabbit |
| RPS6KA (Rsk1-3) P Thr359,Ser363 | CST | 9344 | rabbit |
| S6 | CST | 2217 | rabbit |
| S6 P Ser235,Ser236 | CST | 2211 | rabbit |
| S6 P Ser240,Ser244 | CST | 2215 | rabbit |

| | | | |
|---|---|---|---|
| S6K1 | CST | 9202 | rabbit |
| S6K1 P Thr389 | CST | 9205S | rabbit |
| SLUG (C19G7) | CST | 9585 | rabbit |
| SMAD2 (C86F7) | CST | 3122 | rabbit |
| SMAD2 P Ser465,Ser467 | CST | 3108 | rabbit |
| SMAD2/3 P Ser465/Ser423, Ser467/Ser425 | CST | 8828 | rabbit |
| SMAD3 P Ser423,Ser425 | CST | 9520 | rabbit |
| TCF1 (C63O9) | CST | 2203 | rabbit |
| TCF3 (D15G11) | CST | 2883 | rabbit |
| TCF4 (C9B9) | CST | 2565 | rabbit |
| TGFβ (56E4) | CST | 3709 | rabbit |
| TP53 | CST | 9282 | rabbit |
| TSC2 | CST | 3612 | rabbit |
| TSC2 P Thr1462 | CST | 3617 | rabbit |
| YAP P Ser127 | CST | 4911 | rabbit |
| YAP1 (EP1674Y) | Abcam | ab52771 | rabbit |

**Table S3, Primary antibodies used for RPPA.** CST, Cell Signaling Technology. mAb, monoclonal antibody.

| Target | Accessio n ID (RefSeq) | Fwd primer | Rev primer | Amplic on size (bp) |
|---|---|---|---|---|
| *NANOG* | NM_02486 5.3 | CAGTCTGGACACTGGC TGAA | CTCGCTGATTAGGCTC CAAC | 149 |
| *NODAL* | NM_01805 5.4 | CAGTACAACGCCTATC GCTGT | TGCATGGTTGGTCGGA TGAAA | 75 |
| *POU5F1 (OCT3/4)* | NM_00270 1.5 | TGTACTCCTCGGTCCC TTTC | TCCAGGTTTTCTTTCC CTAGC | 150 |
| *TBP* | NM_00319 4.4 | TAATCCCAAGCGGTTT GC | TAGCTGGAAAACCCAA CTTCT | 170 |

**Table S4. SYBR Green qPCR primers.** Bp, base pairs; Fwd, forward; Rev; reverse. Only a single accession ID is given due to space constraints, but all primers detect additional splice isoforms of their specific target.

**Dataset S1 (separate file).** List of differentially expressed genes in *PIK3CA*$^{WT/H1047R}$ vs wild-type hPSCs after applying an absolute fold-change cut-off of minimum 1.3.

**Dataset S2 (separate file).** List of differentially expressed genes in *PIK3CA*$^{H1047R/H1047R}$ vs wild-type hPSCs after applying an absolute fold-change cut-off of minimum 1.3.

**Dataset S3 (separate file).** List of differentially expressed genes in *PIK3CA*$^{WT/E418K}$ vs wild-type hPSCs.

**Dataset S4 (separate file).** List of differentially expressed genes in *PIK3CA*$^{WT/H1047R}$ vs wild-type MEFs.

**Dataset S5 (separate file).** List of differentially expressed proteins in *PIK3CA*$^{WT/H1047R}$ vs wild-type hPSCs.

**Dataset S6 (separate file).** List of differentially expressed proteins in *PIK3CA*$^{H1047R/H1047R}$ vs wild-type hPSCs.

**Dataset S8 (separate file).** List of differentially expressed transcripts changing in the same direction in both heterozygous and homozygous *PIK3CA*$^{H1047R}$ hPSCs vs wild-type controls.

**Source data deposited on the Open Science Framework**

- High_depth_iPSC_H1047R_RNAseq (related to Fig. 1, 3; Fig. S1, S2)
- M98_iPSCs_E418K_RNaseq (related to Fig. 1)
- MEF_H1047R_RNAseq (related to Fig. 1)
- Total_proteomics (related to Fig. 1, 3; Fig. S2)
- GF_replete_iPSCs_H1047R_RPPA_study (related to Fig. 2; Fig. S3)
- BYL719_iPSCs_H1047R_RPPA_study (related to Fig. 2; Fig. S3)
- WGCNA_iPSC_RNAseq_analysis (related to Fig. 4)
- NODAL_experiment (related to Fig. 5; Fig. S4)
